# Efficient Dilution-to-Extinction isolation of novel virus-host model systems for fastidious heterotrophic bacteria

**DOI:** 10.1101/2020.04.27.064238

**Authors:** Holger H. Buchholz, Michelle Michelsen, Luis M. Bolaños, Emily Browne, Michael J. Allen, Ben Temperton

## Abstract

Microbes and their associated viruses are key drivers of biogeochemical processes in marine and soil biomes. While viruses of phototrophic cyanobacteria are well-represented in model systems, challenges of isolating marine microbial heterotrophs and their viruses have hampered experimental approaches to quantify the importance of viruses in nutrient recycling. A resurgence in cultivation efforts has improved the availability of fastidious bacteria for hypothesis testing, but this has not been matched by similar efforts to cultivate their associated bacteriophages. Here, we describe a high-throughput method for isolating important virus-host systems for fastidious heterotrophic bacteria that couples advances in culturing of hosts with sequential enrichment and isolation of associated phages. Applied to six monthly samples from the Western English Channel, we first isolated one new member of the globally dominant bacterial SAR11 clade and three new members of the methylotrophic bacterial clade OM43. We used these as bait to isolate 117 new phages including the first known siphophage infecting SAR11, and the first isolated phage for OM43. Genomic analyses of 13 novel viruses revealed representatives of three new viral genera, and infection assays showed that the viruses infecting SAR11 have ecotype-specific host-ranges. Similar to the abundant human-associated phage ΦCrAss001, infection dynamics within the majority of isolates suggested either prevalent lysogeny or chronic infection, despite a lack of associated genes; or host phenotypic bistability with lysis putatively maintained within a susceptible subpopulation. Broader representation of important virus-host systems in culture collections and genomic databases will improve both our understanding of virus-host interactions, and accuracy of computational approaches to evaluate ecological patterns from metagenomic data.

## Introduction

It is estimated that viral predation kills ~15% of bacterial cells in marine surface water each day [1] and is a major contributor to nutrient recycling via the viral shunt, where marine viruses make cell-bound nutrients available to the neighbouring microbial community through viral lysis of host cells [2, 3]. Viruses are key players in the modulation of carbon fluxes across the oceans (150 Gt/yr), increasing particle aggregation and sinking to depth [2, 4], and accounting for 89% of the variance in carbon export from surface water to the deep ocean [5]. Viruses alter host metabolism through auxiliary metabolic genes (AMGs), increasing and altering the cellular carbon intake of infected cells [6]. Virus-host interactions also increase co-evolutionary rates of both predator and prey via Red Queen dynamics [7, 8]. While recent metagenomic advances have provided major insight into global viral diversity and abundance [9–12], mechanistic understanding of virus-host interactions in ecologically important taxa is reliant on experimental co-culturing of model systems. In cyanobacteria, such systems have shown that viruses increase duration of photosynthetic function [13] and can inhibit CO2 fixation, providing direct evidence that viruses of abundant phototrophs play an important role in nutrient cycling and global carbon budgets [14]. Furthermore, isolation of new viruses provides complete or near-complete viral genomes, with concrete evidence of known hosts. Such systems are critical to the development, ground-truthing and application of computational methods to identify and classify viral genomes in metagenomic data (e.g. VirSorter [15]; VirFinder [16]; MARVEL [17]); quantify boundaries for viral populations [12, 18] and genera (VConTACT2 [19]); understand the importance of AMGs in altering nutrient flux in natural communities [20] and to predict host ranges of uncultured viruses in metagenomic data (e.g. WIsH [21, 22]).

Viruses of primary producers, such as cyanophages are both well represented with model systems and well-studied in the laboratory. In contrast, virus-host model systems for similarly important and abundant marine heterotrophic bacteria are rare. Isolated viruses infecting heterotrophs are heavily biased towards those with fast-growing, copiotrophic hosts that grow readily on solid agar, enabling the use of plaque assays for viral isolation. Such systems are not representative of the vast majority of heterotrophs in nutrient-limited soil and aquatic environments, which are dominated by slow-growing, oligotrophic taxa with few regulatory mechanisms and complex auxotrophies that limit growth on solid media [23–26]. Advances in Dilution-to-Extinction culturing of ecologically important hosts have enabled the cultivation of many fastidious bacterial taxa that are not amenable to growth on solid media from soil [27], marine [28, 29] and freshwater environments [30]. Without plaque assays to facilitate isolation and purification of viral isolates, cultivation of viruses infecting fastidious taxa in liquid media is challenging, and further exacerbated by the slow growth rates and complex nutrient requirements of their hosts. Paucity of such model systems introduces significant bias in our understanding of viral influence on global carbon biogeochemical cycles. Therefore, it is important that the efforts to isolate heterotrophic bacterial taxa for experimentation and synthetic ecology is matched by efforts to isolate their associated viruses.

Here, we adapted recent advances in Dilution-to-Extinction culturing of hosts [29], and protocols to isolate viruses from liquid media [31] to improve efficiency of cultivating novel virus-host systems for fastidious taxa: First, we selected the ecologically significant SAR11 and OM43 heterotrophic marine clades as models for viral isolation; Second, we used sequential enrichment of viruses from natural communities on target hosts to improve rates of viral isolation [32]; Third, we replaced the requirement for time-intensive epifluorescent microscopy with identification of putative viral infection by comparing infected and uninfected hosts by flow cytometry, followed by confirmation using Transmission Electron Microscopy (TEM). These clades are abundant and important to global carbon biogeochemistry [23, 33–37], but little is known about their associated viruses. In the case of viruses infecting SAR11, two challenges limit our ability to evaluate host-virus ecology in natural communities: (1) Assembly of abundant and microdiverse genomes from viral metagenomes presents a challenge to short-read assembly methods, resulting in underrepresentation in subsequent datasets [12, 18]. This was demonstrated in the successful isolation of the first known pelagiphages by culturing, including the globally dominant HTVC010P, which, prior to its isolation, was entirely missed in marine viromes [38]; (2) poor representation of viral taxa in databases limits our capacity to accurately train machine learning approaches for *in silico* host prediction [12, 39, 40]. This results in either a lack of host information for abundant viral contigs, or worse, incorrect assignment of host to viral contigs, confounding ecological interpretation of data. Some important taxa, such as the OM43 clade, which plays an important role in oxidation of volatile carbon associated with phytoplankton blooms [33, 35, 41], lack any isolated viruses with experimentally confirmed hosts. In addition, both SAR11 and OM43 represent model organisms for genome streamlining as a result of nutrient-limited selection [42]. The effect of genome minimalism on viral infection dynamics is poorly understood, but critical to evaluating the impact of predator-prey dynamics on global marine carbon budgets. In this study, novel SAR11 and OM43 representatives from the Western English Channel were isolated and used as bait to isolate associated viruses. We increased the initial concentration of viruses in natural seawater samples by tangential flow filtration, followed by inoculation of cultures and one to three rounds of sequential enrichment on target hosts in 96-well plates. This yielded 117 viral isolates from 218 inoculated cultures from seven monthly water samples (September 2018 to July 2019). A subsample of putative viral isolates for both clades were sequenced, providing 13 novel viral genomes, including the first known siphovirus to infect SAR11 and the first known virus-host model for OM43.

## Results and Discussion

### Isolation of a novel SAR11 strain and three new OM43 strains from the Western English Channel to use as bait for phage isolation

Dilution-to-Extinction culturing for host taxa using natural seawater-based medium was performed from a water sample collected in September 2017 from the Western English Channel, and yielded the first SAR11 strain (named H2P3α) and the first three OM43 strains (named C6P1, D12P1, H5P1) from this region. The full-length 16S rRNA gene of *Pelagibacter sp*. H2P3α was 100% identical to that of the warm water SAR11 ecotype *Pelagibacter bermudensis* HTCC7211 (subclade 1a.3) and was considered to be a local variant [43] (Supplementary Fig. 1A). All three novel OM43 isolates were most closely related to *Methylophilales sp*. HTCC2181, a streamlined member of the OM43 clade with a 1.3 Mbp genome, isolated from surface water of the North Eastern Pacific [44] (C6P1 96.17%, D12P1 96.62%, H5P1 97.79% nucleotide identity across the full 16S rRNA gene) (Supplementary Fig. 1B). The average nucleotide identity of the 16S rRNA gene of isolates CP61, D12P1 and H5P1 to each other was ~98.46% (Supplementary Table 1), suggesting they are representatives of the same genus [45].

### An efficient, low-cost method of isolating new viruses yielded 117 new viral isolates for SAR11 and OM43 taxa

Using the four new hosts from above and established SAR11 isolates *Pelagibacter ubique* HTCC1062 (subclade 1a.1) and HTCC7211 (subclade 1a.3), we developed an optimised viral isolation pipeline (Fig. 1) and applied it to six monthly water samples from the Western English Channel, taken between September 2018 to April 2019 (Supplementary Table 2). Each month, we concentrated a natural viral community with tangential flow filtration and used it to inoculate one to two 96-well Teflon plates containing host cultures at ~10^6^ cells·mL^-1^. Plates were monitored by flow cytometry and growth of putatively infected cultures was compared to those of unamended controls over the course of ~2 weeks to account for the slow growth rates of SAR11 and OM43 [35, 46]. An additional sample was taken in July 2019 to attempt viral isolation on OM43 strains D12P1 and H5P1. Out of a total of 218 cultures amended with concentrated viral populations, 117 viruses were isolated, purified and still infective after at least three passages, with repeatable differences observed in cytograms of infected cultures and control cultures (Supplementary Fig. 2-5). This represents an overall isolation efficiency of 53% and an average yield of 18 viruses per environmental sample (Fig. 2). For 90% of inoculated SAR11 cultures (94 out of 105) we observed evidence of viral infection, and the putative viral isolate could be propagated and purified, fulfilling Koch’s postulates for confirming a pathogenic agent. For OM43, 23 out of 113 (20%) inoculated cultures yielded positive infections that could be similarly propagated. All viral isolations required between one and three rounds of virus enrichment (Supplementary Table 3) before changes in host growth curves between infected and uninfected cultures could be observed. This suggests putatively rare viruses can be enriched within three rounds to a level at which infection can be observed on a flow cytometer.

**Fig. 1.**
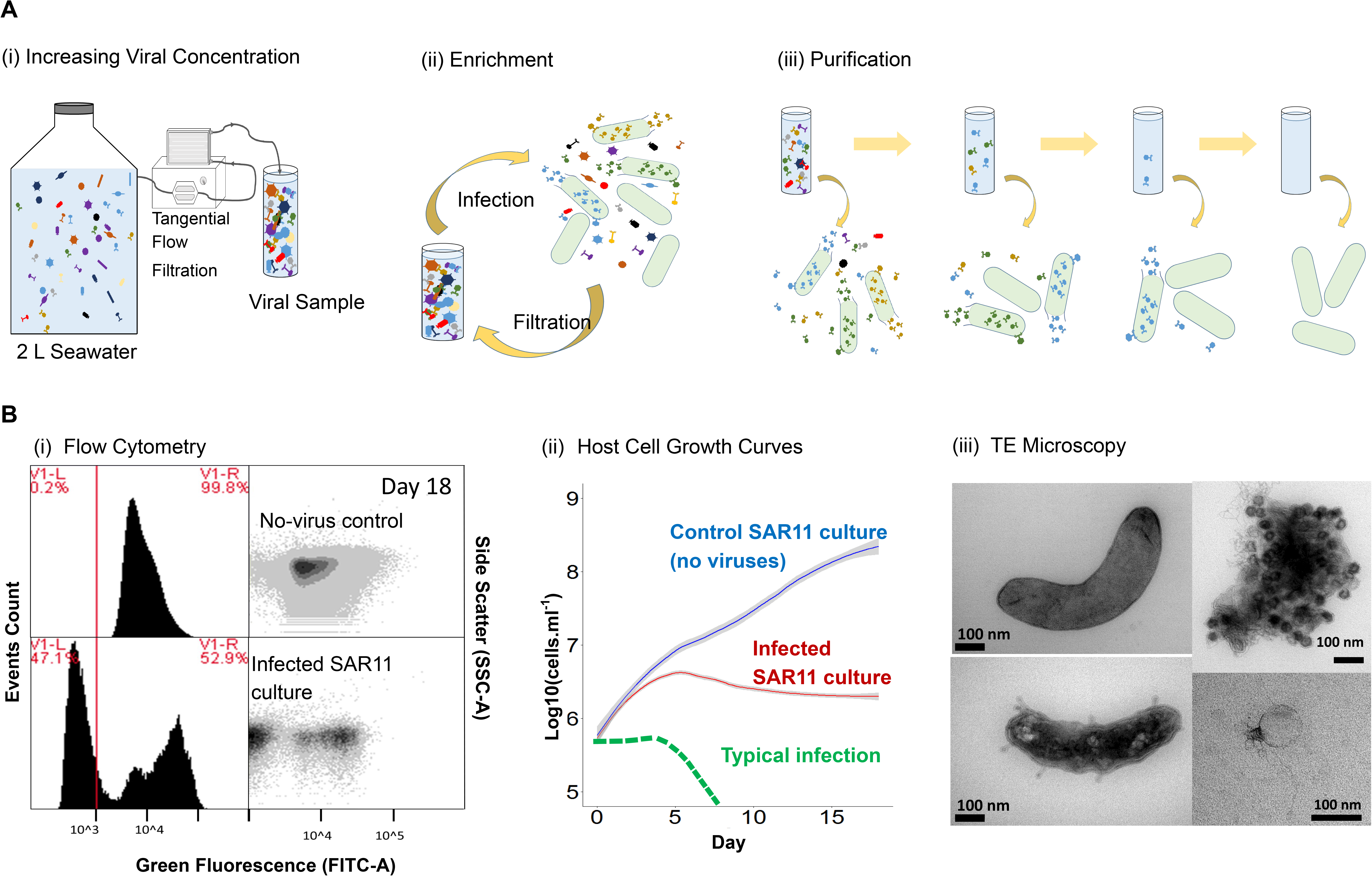
Workflow for high-throughput isolation **(A)** i. Increasing concentration of viruses in water samples by tangential flow filtration (TFF); ii. Initial infection of host cultures to enrich the sample for specific viruses; iii. Purification of viral isolates through three rounds of Dilution-to-Extinction; **(B)** Initial screen of viral infections using (i). Flow cytometry, by comparing populations of no-virus controls and infected cultures; (ii). Comparing growth curves of no-virus control culture (HTCC1062) against infected SAR11 cultures; (iii). Confirming the presence of viruses in infected SAR11 cultures using transmission electron microscopy: Top left: HTCC1062 no-virus control, bottom left: infected HTCC1062, top right: aggregated cellular debris and viruses, bottom right: virus found in infected HTCC1062 culture.

**Fig. 2.**
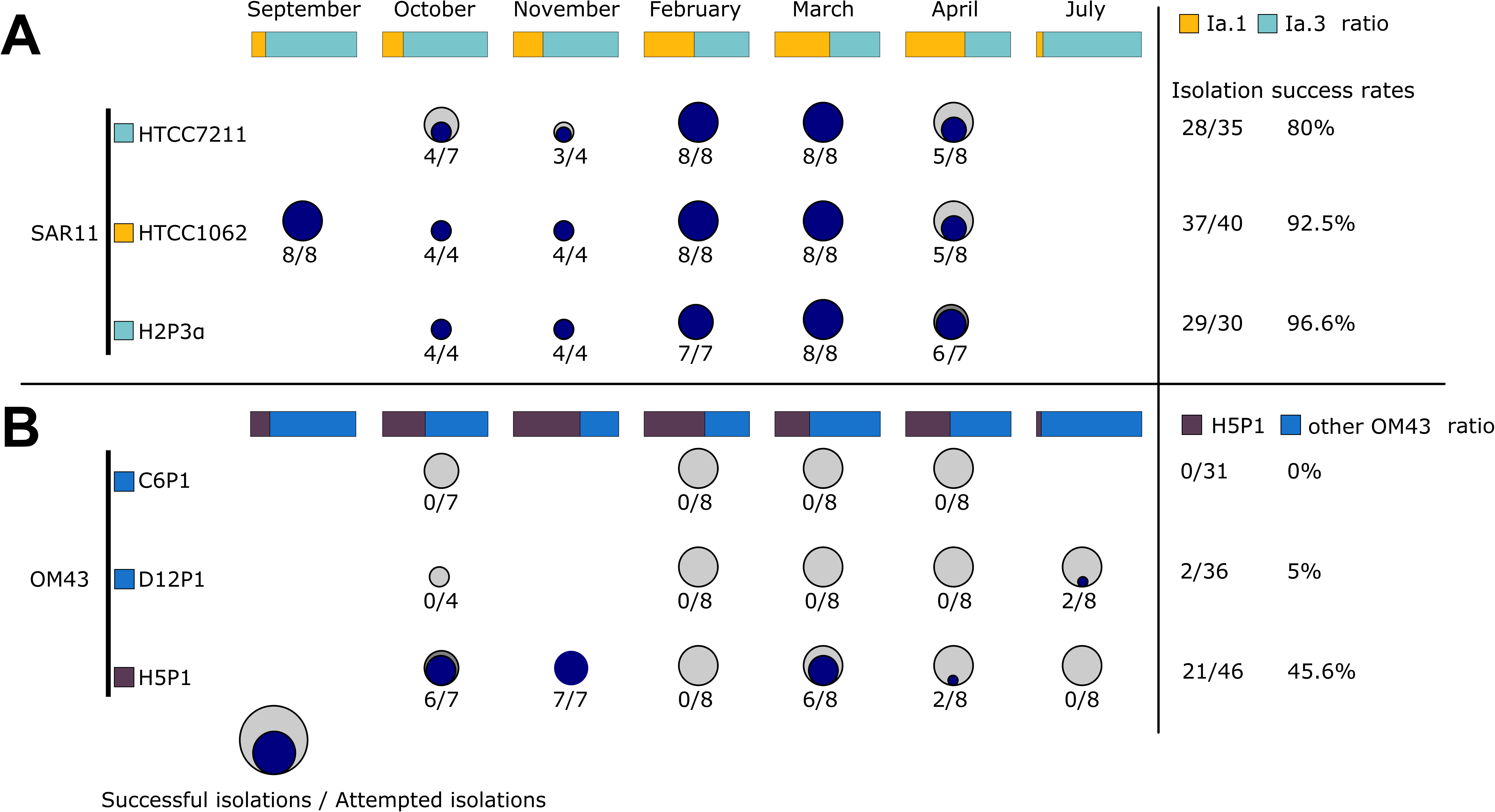
Summary of success rates for each bacterial host as well as relative contribution to the clade community during sampling based on 16S rRNA analysis: (**A**) Three SAR11 strains HTCC1062, HTCC7211 and H2P3α and (**B**) three OM43 strains C6P1, D12P1 and H5P1. Circles and associated numbers represent number of successful isolations (i.e. successful passage through three rounds of Dilution-to-Extinction), over the total number of attempts made. Bars at the top of each panel represent the relative abundance (SAR11 subclades 1a.1 and 1a.3, and OM43 strains for A and B, respectively) of strains in the microbial community sampled at the time of collection (based on 16S rRNA community profiling). Percentages on the right represent total successes for all attempts per host strain.

### New viruses represent novel viral populations and support established ANI cut-offs for ecologically discrete viral ecotypes

Due to the rate-limiting step of culturing sufficient biomass for extraction of viral DNA, we subselected 16 viral isolates based on availability in November 2018 across four different hosts (HTCC1062, HTCC7211, H2P3α and H5P1) for Illumina sequencing to >30-fold coverage (Table 1). Three out of 16 sequenced samples (~19%, two from host HTCC7211, one from OM43 host H5P1) failed to assemble into single viral contigs, in line with previously reported failure rates of 18-39% for phages of *Escherichia coli* and *Salmonella spp*. [47]. For 11 of the remaining 13 samples (12 from SAR11 hosts and one from OM43), each individual sequence assembly was identified as a complete viral genome by VirSorter [15] and 95-100% complete using CheckV [48]. All assemblies yielded a single viral contig (categories 1 or 2, >15kbp) per sample, indicating that our purification process was effective in recovering pure viral isolates. Interestingly, all viral isolate genomes were classified by VirSorter as Category 2 (“likely virus”) rather than Category 1 (“most confident”), despite being complete - indicating either a lack of viral hallmark genes or a lack of enrichment of viral genes on the contigs. This finding matches our own observations for other isolated viruses infecting SAR11 (data not shown), and suggests VirSorter classification of pelagiphages is conservative. Viral isolates were named in accordance with current ICTV guidelines for prokaryotic virus taxonomy [49], with names selected from Norse mythology and folklore, and contemporary culture (Table 1).

**Table 1.**
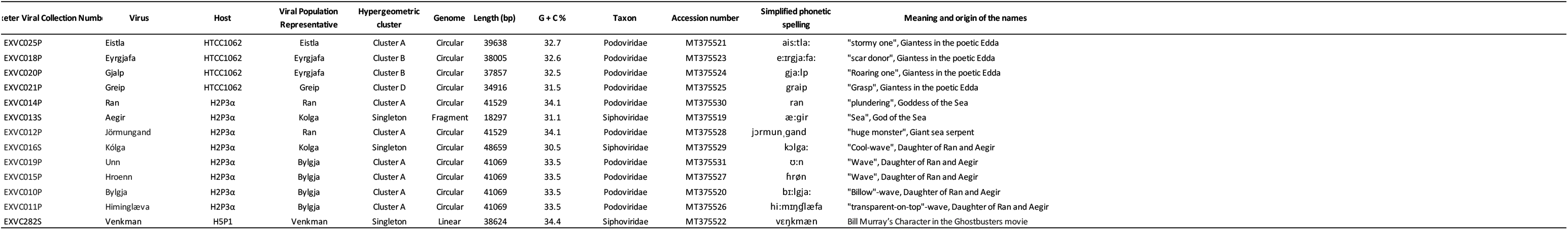
Summary of phages isolated and sequenced in this study.

Viral populations are defined as discrete ecological and evolutionary units that form non-overlapping clouds of sequence space, and previous work in cyanophage populations have shown that viral populations can be delineated into populations using an average nucleotide identity (ANI) cutoff of 95% [50]. Pairwise ANI was calculated between the thirteen successfully sequenced viral genomes from this study and 85 other known or putative pelagiphages [39, 51, 52]. Pairwise ANI ranged between 77.5-100%, with a discrete distribution between 96.4-100.0% (Supplementary Fig. 6, Supplementary Table 4). This is in agreement with previous work in cyanophages [50, 53] and supports the broad use of proposed boundary cutoffs to define viral populations within viral metagenomic assemblies [10, 11]. At the proposed ANI cutoffs of 95% over 85% length [18], our 13 new viruses clustered into six viral populations, ranging from singletons to a viral population with four members (Table 1). Phages within the same populations were all isolated from the same environmental sample and on the same host, in agreement with their classification as discrete ecological and evolutionary units. All viruses isolated in this study formed their own exclusive viral populations, with no representatives from either known isolates [38, 52] or fosmid-derived [39] genomes from other studies, indicating that a high degree of viral population diversity remains to be discovered in the Western English Channel and beyond.

### Current pelagiphage isolates can be organised into five distinct phylogenetic clades

To evaluate isolate diversity at higher taxonomic organisation, we picked one representative from each of our six viral populations and compared them to previous isolates and fosmid-derived phage sequences using three approaches: First, phylogenetic analyses were performed based on conserved genes in known pelagiphage isolates and closely-related taxa from viral metagenomic surveys [11, 12]; Second, raw hypergeometric probability of shared gene content was calculated (to capture broader relationships and account for genomic mosaicism) [54]; Third, genomes were organised into ICTV-recognized genera using vConTACT2. vConTACT2 initially derives viral clusters using a hypergeometric approach, with subsequent refinement with Euclidean-distance-based hierarchical clustering to split mismatched, ‘lumped’ viral clusters [19]. All three approaches were congruent - clustering on probability of shared gene content organised pelagiphage genomes into four main clusters and numerous singleton genomes (Fig. 3). This was broadly supported by phylogenetic (Supplementary Fig. 7) and vConTACT2 (Supplementary Fig. 8) classification approaches. Cluster A contained 23 members (nine from fosmid-derived contigs [39]; eleven previously isolated pelagiphages [52, 55]) and Pelagibacter phages *Ran* (EXVC014P), *Bylgja* (EXVC010P) and *Eistla* (EXVC025P) from this study. Cluster B contained two previously isolated pelagiphages, one fosmid-derived contig and Pelagibacter phage *Eyrgjafa* (EXVC018P) from this study. All viruses in Clusters A and B were assigned to a single viral genus by vConTACT2 that also contained 12 previously isolated pelagiphages [38, 52]. Cluster C only contained fosmid-derived contigs from the Mediterranean [39], with no isolated representatives, marking it an important target for future isolation attempts. Cluster D contained eight fosmid-derived contigs, Pelagibacter phage HTVC010P, and Pelagibacter phage *Greip* (EXVC021P) from this study. Pelagibacter phage *Kolga* (EXVC016S) from this study and the only known Pelagimyophage HTVC008M from a previous study [38] fell outside the four main clusters. VConTACT2 split Cluster D, leaving Pelagibacter phages *Greip* and *Kolga* as members of two singleton clusters, suggesting they are the first cultured representatives of novel viral genera and distinct from the globally ubiquitous Pelagibacter phage HTVC010P.

**Fig. 3.**
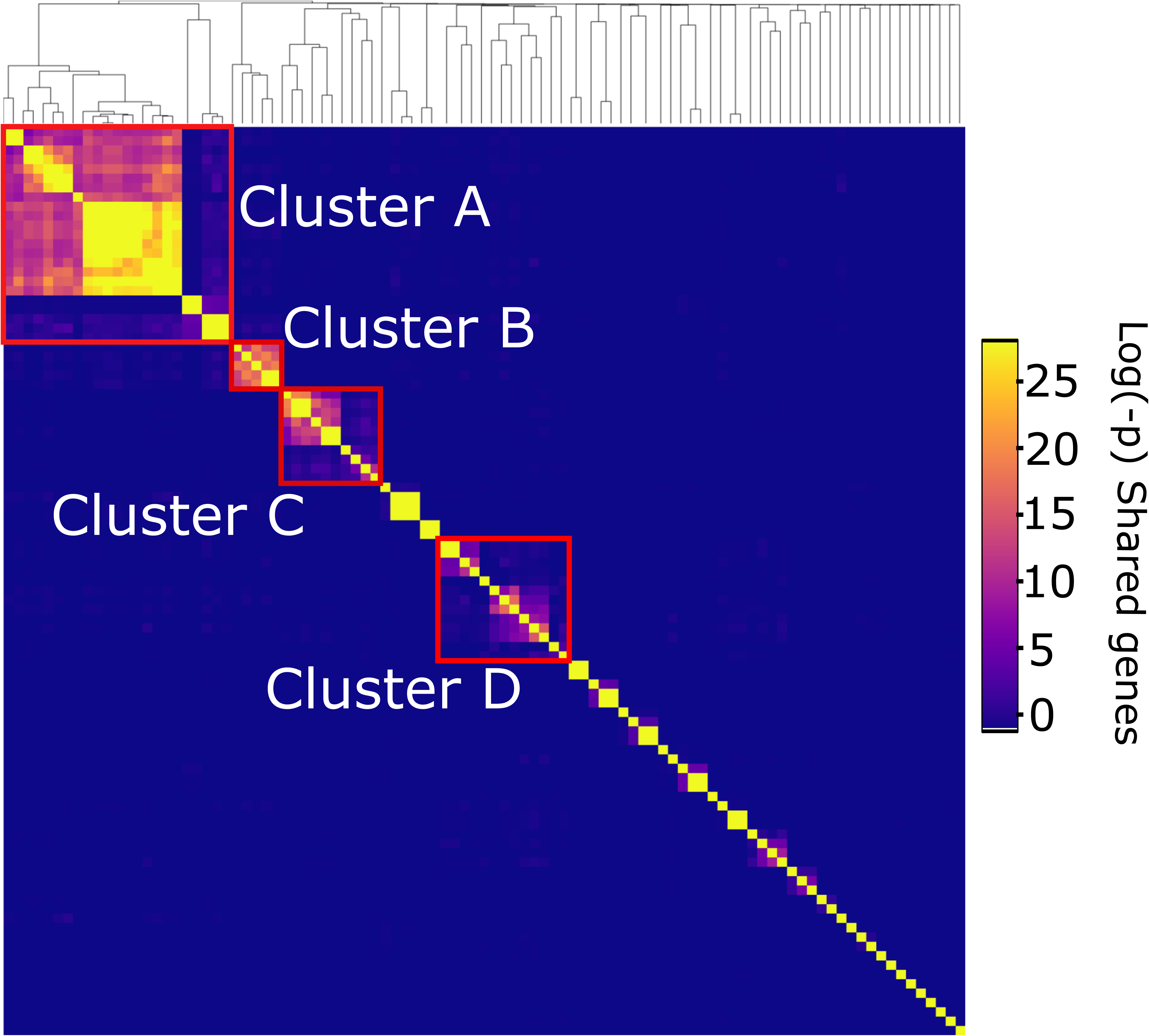
Hypergeometric probability of shared gene content between known pelagiphage genomes identified four main viral phylogenetic clusters (outlined with red boxes).

### Novel pelagiphages are ecotype-specific and persist in the community

Community composition analysis using 16S rRNA genes showed that during the sampling period (September 2018 to July 2019) the SAR11 contribution relative to the total number of sequences ranged from a minimum of 28.0% in July 2019 to a maximum of 42.2% in September 2018 (Supplementary Fig.9). At the clade level, SAR11 composition was relatively stable over time, except for clade II roughly doubling its relative contribution from 15.1% to about 29.6% between February and April. The SAR11 community was dominated by clade I overall throughout the sampling period. Within clade I, the ratio between SAR11 subclade Ia.1 (cold-water ecotype) and Ia.3 (warm-water ecotype) showed that the warm-water ecotype dominated from September to November as well as July (Figure 2A). During the coldest months in the Western English Channel (February to April) the cold-water ecotype became as abundant as the warm-water ecotype with roughly a 1:1 ratio. Overall, temperature ranged from 9.3 °C - 14.1 °C from October to April, when isolations were attempted on all strains (Supplementary Table 2). HTCC1062 and HTCC7211 have specific growth rates of ~0.22 and ~0.12 divisions per day at 10 °C, respectively [46]. Our new SAR11 isolate from the Western English Channel, *Pelagibacter sp*. H2P3α, showed similar growth rates to *Pelagibacter bermudensis* HTCC7211 (Supplementary Fig. 10), with specific growth rates of ~0.10, ~0.45 and ~0.84 divisions per day at 10, 18 and 25 °C, respectively. Therefore, measured *in situ* temperatures during our sampling period were sufficient to support slow growth of warm-water ecotypes even during winter months, potentially providing sufficient prey to support a population of warm-water ecotype specific phages. This result shows that slow growth of warm water ecotypes at *in situ* temperatures is possible, supporting the finding of the 16S community analysis that this ecotype persists throughout the year.

An alternative explanation is that isolated viruses have a broad host range that encompasses both warm- and cold-water ecotypes of the SAR11 subclade Ia. We tested the host range (Table 2) of six pelagiphages (one of each viral population), isolated from samples in October 2018 and November 2018 (*in situ* water temperature of 14.8 °C and 14.2 °C respectively) across the three SAR11 strains (Table 1). Pelagibacter phages *Eistla, Eyrgjafa* and *Greip* all infected cold-water ecotype HTCC1062 exclusively, while Pelagibacter phages *Ran* and *Kolga* only infected warm-water ecotypes HTCC7211 and H2P3α. Pelagibacter phage *Bylgia* was the only virus that could infect both warm and cold-water ecotypes. Therefore, our new pelagiphages appear to be broadly ecotype-specific, confirming previous findings [52]. Our results suggest overall that pelagiphages persist in the water column throughout the year in sufficient densities to be isolated by our enrichment method, despite ecotypic specificity and fluctuations in warm- and cold-water ecotype community contributions of SAR11 subclade 1a. If concentration and enrichment of viruses during isolation is sufficient to successfully isolate even low abundance phages then a comprehensive library of representative phage isolates could be generated with relatively modest sampling effort across a few locations.

**Table 2.** Host infectivity of viral populations isolated and sequenced in this study. *Pelagibacter sp*. H2P3α and *Pelagibacter bermudensis* HTCC7211 are warm-water ecotypes of SAR11 subclade 1a; HTCC1062 is a cold-water ecotype*. Pelagibacter* phage *Bylgja* was the only virus capable of infecting both ecotypes.

| Phage | H2P3 | HTVC1062 | HTVC7211 |
| --- | --- | --- | --- |
| Eistla |  | + |  |
| Eyrgjafa |  | + |  |
| Greip |  | + |  |
| Ran | + |  | + |
| Kolga | + |  | + |
| Bylgja | + | + | + |

### Pelagibacter phages Kolga and Aegir-the first siphovirus infecting SAR11

The 25 previously known viral isolates infecting SAR11 comprise 24 podoviruses and one myovirus [38, 52, 56]. Previous cultivation efforts for viruses of SAR11 have not isolated any siphoviruses, nor are any known from viral metagenomic studies. We isolated and sequenced the two Pelagibacter phages *Kolga* and *Aegir* using host H2P3α as bait (Fig. 4A-C). Transmission electron microscopy (TEM), which showed evidence of a long tail (Fig. 4B), suggested classification as the first reported siphoviruses infecting members of the SAR11 clade.

**Fig. 4.**
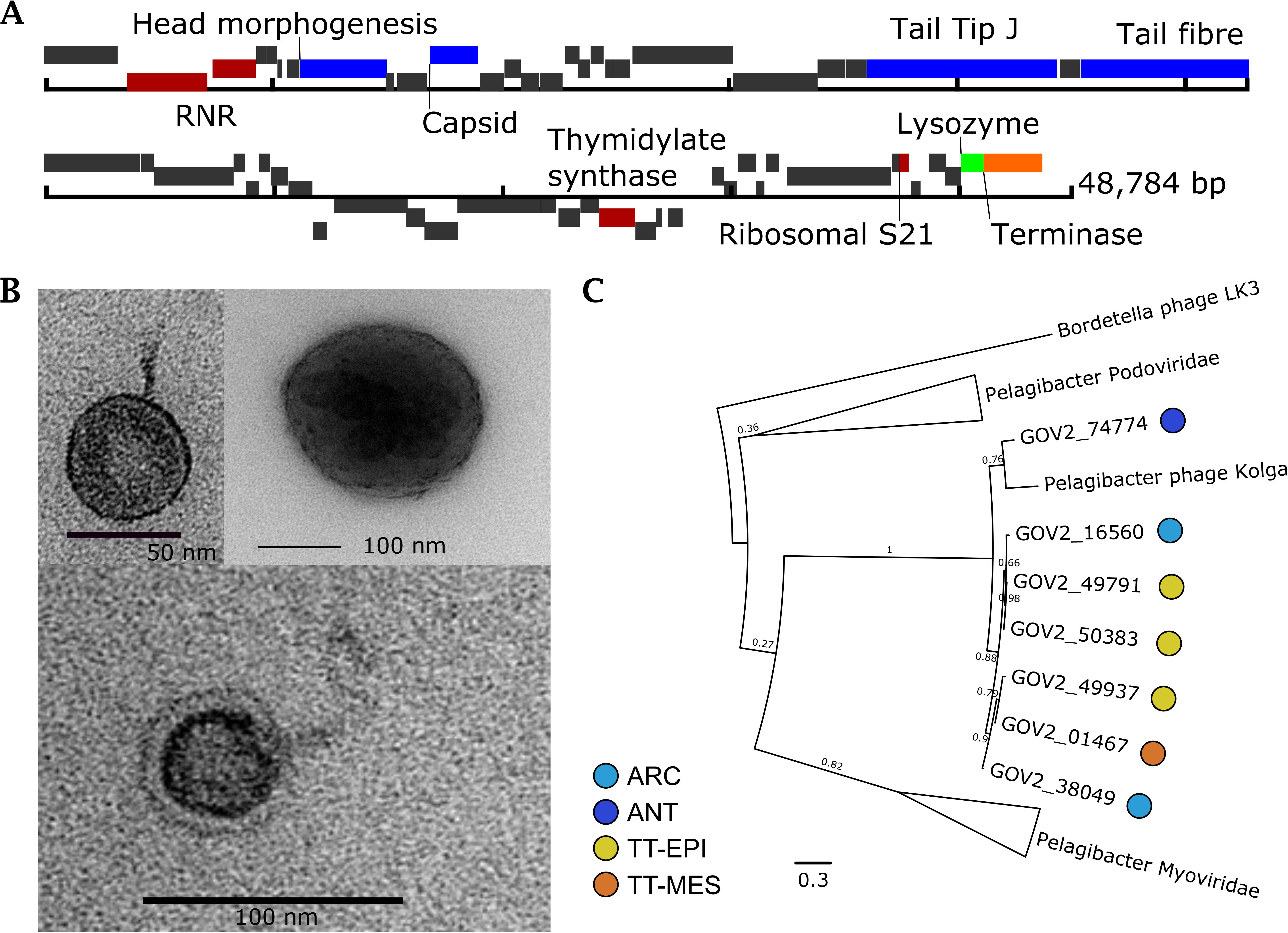
The first reported siphovirus infecting SAR11 (Pelagibacter phage *Kolga* EXVC016S) isolated on novel host *Pelagibacter sp*. H2P3α. **(A)** Gene map of the 48,784 bp genome, which contains 80% hypothetical genes without known function; **(B)** Transmission Electron Micrographs of *Kolga* (left and bottom) and an H2P3α host cell infected with *Kolga* (top right). **(C)** Unrooted maximum likelihood tree of *terL* genes found in pelagiphages. Branches containing members of the Podoviridae and Myoviridae family infecting SAR11 are collapsed for clarity (full tree is available in Supplementary Fig. 7). Closely related contigs prefixed with ‘GOV’ represent viral contigs from the Global Ocean Virome dataset and the ecological zone from which they were assembled is marked: Temperate-tropical epipelagic (TT-EPI); temperate-tropical mesopelagic (TT-MES); Antarctic (ANT), Arctic Ocean (ARC).

Pelagibacter phages *Aegir* and *Kolga* were classified as members of the same population using a boundary cutoff of 95% ANI over 85% contig length, however, *Aegir* had a length of 18,297 bp compared to 48,659 bp in *Kolga*, therefore we considered *Aegir* to be a partial genome of the same viral population. *Kolga* did not share a significant number of genes with known SAR11 podoviruses (Fig. 3), and did not cluster with other pelagiphages using hypergeometric analysis based on shared gene content. VConTACT2 also grouped *Aegir* and *Kolga* into one cluster without any other known viruses, suggesting it represents a novel viral genus. A number of genes found in *Kolga* were shared with other known siphoviruses (associated with different hosts) such as the Bordetella phage LK3. Screening of contigs from the Global Ocean Virome dataset [11] identified six contigs from various ecological zones which shared a viral cluster with *Kolga*, but which belonged to different viral populations based on network analysis using vConTACT2. Phylogenetic analysis of all Pelagibacter phage TerL genes indicates that the closest known relatives to *Kolga* and *Kolga*-like contigs are members of the Myoviridae family, supporting a classification as a distinct and novel viral group (Fig. 4C, Supplementary Fig. 7). In *Kolga* 67% of encoded genes could not be functionally annotated, and out of all hypothetical genes identified on *Kolga*, only three hypothetical genes were shared with SAR11 podoviruses. In contrast, on average ~90% of genes without known function identified within our novel SAR11 podoviruses are shared between different SAR11 podoviruses. *Kolga* possesses a tail tip J protein (Fig. 4A), often found in phages with long non-contractile tails such as E. coli phage λ, where it plays a role in DNA injection during cell entry and tail assembly [57]. *Kolga* also encodes a small S21 subunit of the 30S ribosomal gene structurally similar to the ones found in Pelagibacter phage HTVC008M, Pelagimyophage-like contigs [58], and hosts HTCC7211 and H2P3α. Encoding ribosomal genes is a feature found in numerous myoviruses and siphoviruses [59]. The S21 gene is involved in translation initiation and needed for the binding of mRNA [60]. Virally-encoded S21 genes may provide a competitive advantage for the phage as it could replace cellular S21 and assist in the translation of viral transcripts. *Kolga* may need its S21 gene for shifting the translational frame, as it has been shown that for some members of the Caudovirales the production of tail components is dependent on programmed translational frameshifting [61]. Given the constitutive nature of gene expression in genomically streamlined bacteria [62], genes such as S21 may also provide the virus with a mechanism to manipulate host metabolism in the absence of typical promoters and repressors.

### First methylophages for marine OM43 isolated

Isolation of novel phages for the OM43 clade yielded 23 positive infections, with efficiencies ranging from 0% (no viruses isolated on host C6P1) to 45% on H5P1 (Fig. 2). To the best of our knowledge these are the first reported viruses infecting members of the OM43 clade. One explanation for the lower efficiency of isolation of phages infecting OM43 is simply one of lower host abundance concomitant with lower phage abundance in the viral community, reducing the likelihood of infective viruses coming into contact with susceptible and permissive cells. OM43 are closely associated with metabolism of extracellular substrates from phytoplankton blooms [63], but have low abundance outside of phytoplankton spring blooms [33]. Our water samples were not associated with high *insitu* fluorescence (used as proxy measurement for phytoplankton), and missed the April 2019 spring bloom by about two weeks (Supplementary Fig. 11). Based on 16S community analysis, the OM43 contribution to the bacterial community (0.7%) was lowest during sampling in April 2019 and highest in October 2018 (1.9%). Relative to all other OM43, H5P1 was the most abundant OM43 in the Western English Channel, contributing more than half of all OM43 in November (Fig. 2B). This could explain the higher success rate of isolating phages on host H5P1 compared to C6P1 and D12P1. Sequencing of the first OM43 phages isolate, Methylophilales phage *Venkman* (EXVC282S), returned a single genome 38,624 bp long (31.9% GC content), which was linear but complete (Supplementary Table 5). *Venkman* encodes genes (Fig. 5A) with similar synteny and function to the siphovirus P19250A (38,562 bp) that infects freshwater *Methylophilales* LD28, which are often considered a freshwater variant of OM43 [64, 65]. Unlike the siphovirus P19250A, TEM images indicated that OM43 phage *Venkman* had a short tail (Fig. 5B) similar to podoviruses, though it is possible that tail structures were lost during grid preparation. *Venkman* shared a viral population with a 23kbp contig from the Western English Channel virome assembled from short-read data (WEC_HYBRID_01170), suggesting this viral type does not suffer from the issues of high abundance and microdiversity that challenge assembly of some pelagiphages. Phylogenetic analysis of concatenated TerL and exonuclease genes indicated that *Venkman* is most closely related to P19250A and other siphoviruses infecting different Proteobacteria (Fig. 5C). However, branch support values were low, despite numerous attempts to refine the tree with different approaches (see Supplementary Methods). We identified a number of common phage proteins such as a capsid protein, terminase, nucleases and tail structural proteins, and the remaining 54% of genes were hypothetical. VConTACT2 assigned OM43 phage *Venkman* and LD28 phage P19250A to the same genus-level cluster, therefore the OM43 phage *Venkman* may be a marine variant of the freshwater LD28 siphovirus P19250A.

**Fig. 5.**
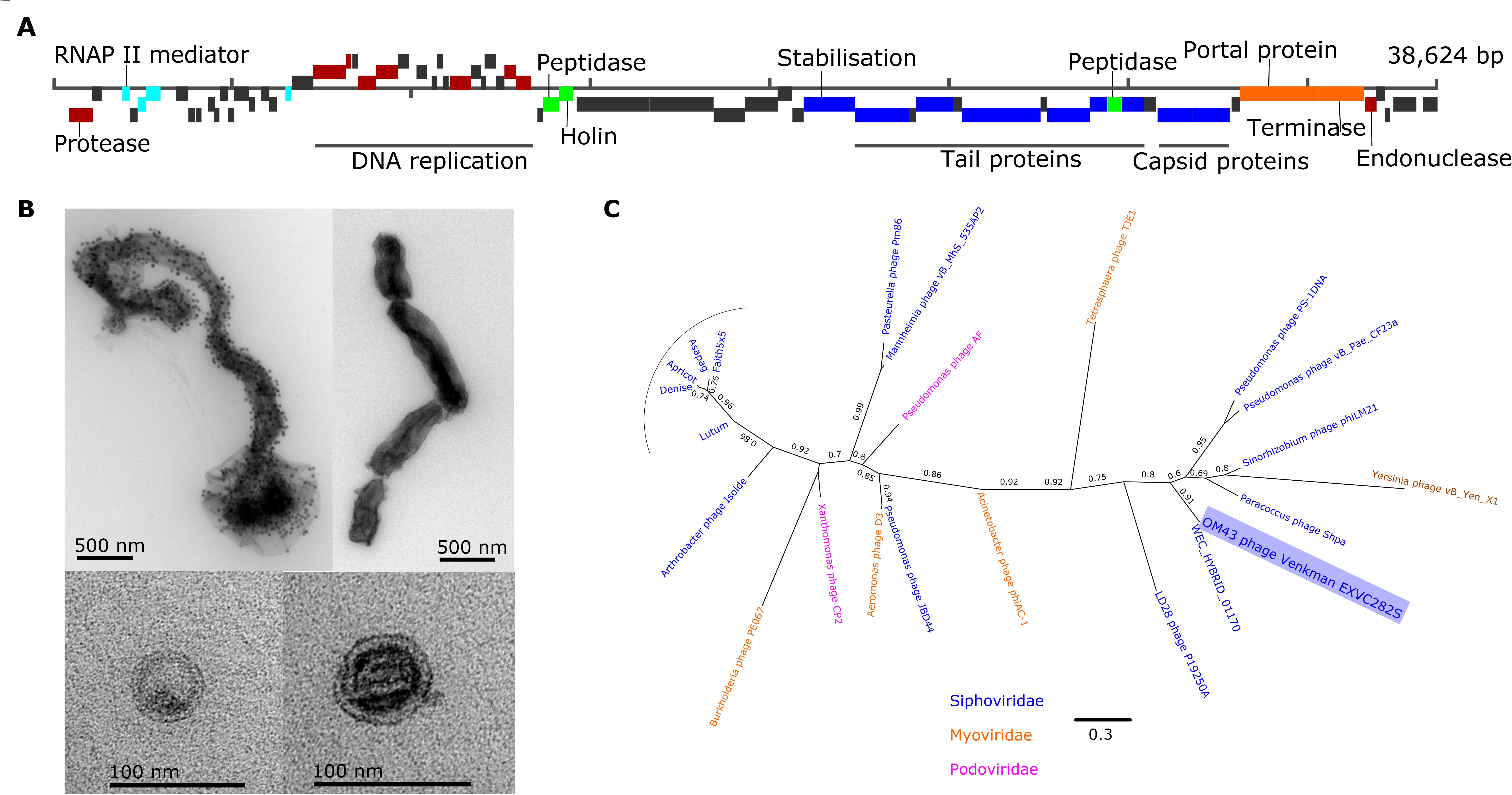
The first reported cultured virus known to infect a member of the OM43 clade (Methylophilales phage *Venkman* EXVC282S) which was isolated on host strain H5P1 from this study; **(A)** Gene map displaying protein coding genes. Gene function is colour coded as follows: Structural genes (blue); DNA replication (red); lysis (green); transcription (turquoise); packaging (orange); hypothetical genes (grey). **(B)** TEM images of infected and chaining H5P1 cells (top left), uninfected H5P1 chaining cells (top right), *Venkman* viral particles (bottom left and right); **(C)** Maximum likelihood tree (500 bootstraps) of concatenated viral *terL* and exonuclease genes. Host families of the phages are indicated on the figure.

### Global abundance of novel isolates highlights niche-specificity and low representation in existing datasets

To evaluate relative global abundance of our new phage isolates, existing virome datasets from the Global Ocean Virome survey [11] and the Western English Channel [12] were randomly subsampled to 5 million reads and competitively recruited against genomes of viral population representatives from this study as well as previously isolated pelagiphages (see Supplementary methods). Overall, with the exception of Pelagibacter phage HTVC010P, pelagiphages were poorly represented in samples from temperate-tropical mesopelagic (TT-MES) and Antarctic (ANT) ecological zones, as defined in [11] (Fig. 6). Phages isolated in this study were neither globally ubiquitous, nor abundant in the single Western English Channel virome (with the exception of phage *Ran*). Phages *Bylgja, Himinglaeva, Eistla* and *Eyrgjafa* did not achieve the minimum cutoff of 40% genome coverage to be classified as present [18] in any of the viromes tested. *Ran* was the only phage isolated in this study with representation in at least two samples from temperate-tropical epipelagic (TT-EPI) zones, concordant with its host specificity of warm-water SAR11 ecotypes (*Pelagibacter bermudensis* HTCC7211 and H2P3α from this study (Table 2). Similarly, the Western English Channel virome was taken in September 2016, when waters are usually highly stratified after summer heating [66] and warm water ecotypes of SAR11 dominate the microbial community (Fig. 2A). Other viral populations from this study also isolated on warm water SAR11 ecotypes (phages *Kolga, Bylgja* and *Himinglaeva*) were either absent or below limits of detection in this sample. This coupled with the low global abundance of these viruses suggests that pelagiphage communities comprise few highly abundant taxa and a long rare tail. Pelagibacter phage *Greip* was detected in seven samples, six of which were Arctic (ARC) samples from a discrete region of the Arctic characterized by low nutrient ratios [11]. In three of those samples (191_SRF, 193_SRF, 196_SRF), HTVC010P was not detected and in the other three (206_SRF, 208_SRF and 209_SRF) *Greip* was 1.5 to 6.6-fold more abundant than HTVC010P, identifying *Greip* as an abundant arctic pelagiphage ecotype. The fact that *Greip* was isolated on the cold-water ecotype of SAR11 (Pelagibacter ubique HTCC1062) and does not infect either warm-water ecotype HTCC7211 or H2P3 supports the hypothesis that host niche specificity shapes the phylogeography of associated viral taxa.

**Fig. 6.**
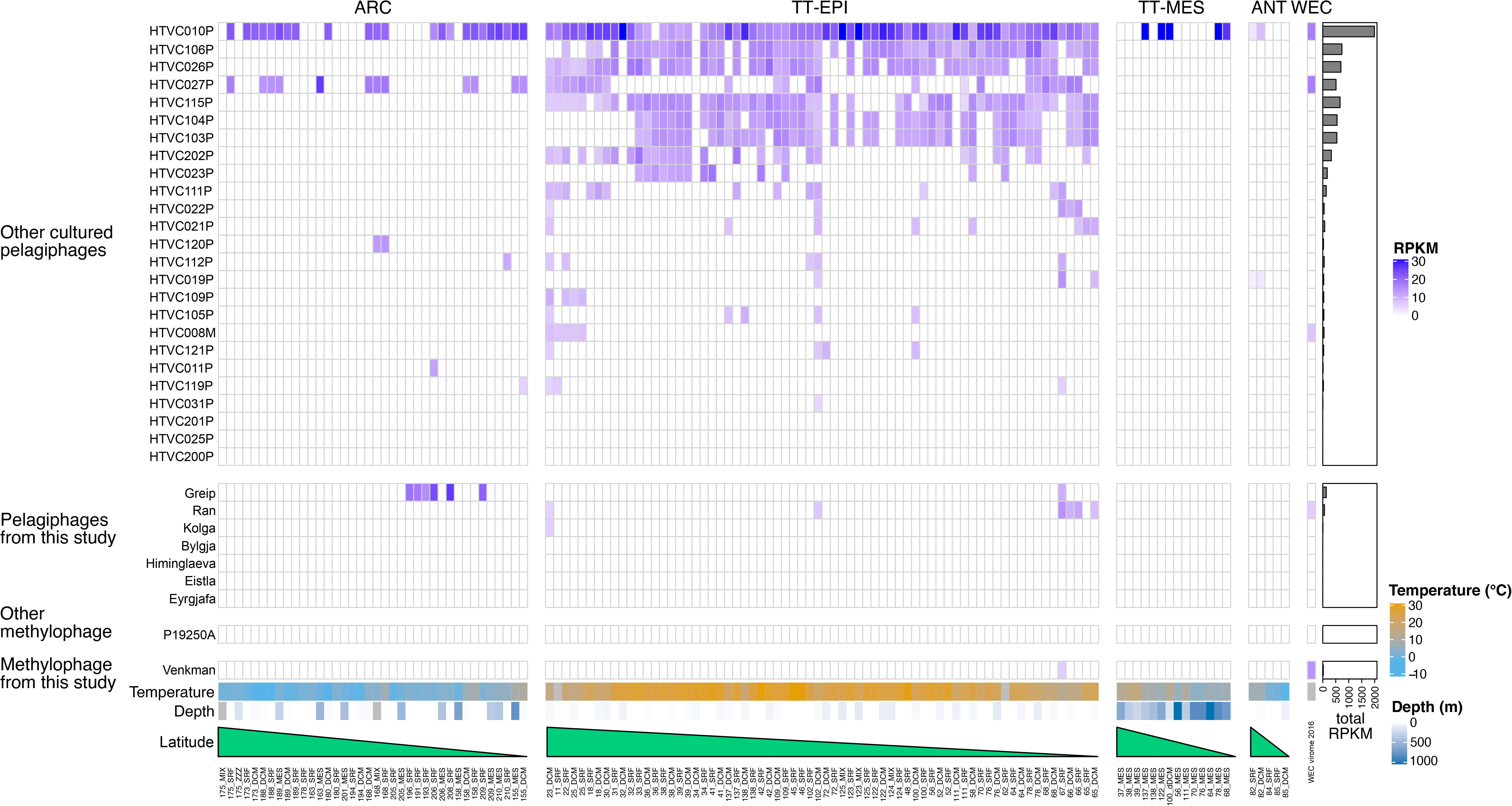
Relative abundance (reads recruited per kilobase of contig per million reads (RPKM)) of known pelagiphage isolates and methylophage genomes in viromes from the Western English Channel [12] and the Global Ocean Virome [11]. Samples are organised by decreasing latitude per ecological zone: ARC: Arctic; TT-EPI: temperate-tropical epipelagic; TT-MES: temperate-tropical mesopelagic; ANT: Antarctic; WEC: Western English Channel.

In the Western English Channel viral metagenomes, OM43 phage *Venkman* was the third most abundant (2,699 RPKM) after Pelagibacter phages HTVC027P (5,023 RPKM) and HTVC010P (5,026 RPKM) identifying it as an ecologically important virus in this coastal microbial community (Fig. 6). In contrast, neither *Venkman* nor LD28 phage P19250A were identified in the majority of samples from GOV2, except in one sample (67_SRF) in the South Atlantic Ocean (257 RPKM against *Venkman*). Sample 67_SRF was also classified as a coastal biome by the authors [11]. These results coincide with previous reports that OM43, and presumably their associated viruses as a consequence of host abundance, are important in some coastal regions, but largely absent in open-ocean systems [33, 41].

### Unusual host-virus dynamics are prevalent in isolated phages

Pelagibacter phages *Bylgja, Eyrgjafa, Ran* and *Eistla* all encode endonucleases and exonucleases (Supplementary Fig. 12) and cluster by shared protein content with other pelagiphages (Cluster A and B), such as e.g. HTVC011P and HTVC025P, shown previously to integrate into host genomes (Fig. 3) [52]. We therefore predict that all our phages within Clusters A and B are temperate phages. *Eyrgjafa* also encodes a tRNA-Leu, that has 85% nucleotide identity over the first 34 bases of the tRNA-Leu of its host HTCC1062, suggesting a putative integration site into the host genome [67]. To date, 16 of the 29 viruses previously isolated on SAR11 strains have either been shown to be capable of lysogeny, and/or encode genes associated with a temperate infection cycle. In contrast, viruses such as the Pelagibacter phages *Greip*, and the abundant HTVC010P in Cluster D do not possess any genes associated with lysogeny, and would therefore be classified as exclusively lytic. Viruses in Clusters A and B (putatively temperate phage), were of much lower abundance in the environment compared HTVC010P which was among the most abundant pelagiphages in the Western English Channel (Fig. 6), suggesting a possible ecological difference between two groups of viruses with different infectivity strategies.

Interestingly, growth curves of hosts infected with Pelagibacter phage *Greip* and other isolated phages deviated from the expected decay in cell abundance associated with viral lysis and previously observed in isolated pelagiphages [38] (Fig.1 Bii). The first pelagiphages HTVC010P and HTVC008M were isolated from the warm waters of the Sargasso Sea, and HTVC011P and HTVC019P were isolated from the colder waters of the Oregon Coast. All four strains were propagated on the cold-water SAR11 ecotype *P. ubique* HTCC1062. In all cases, host density was reduced from ~8 × 10^6^ cells·mL^-1^ at T_0_ to <10^6^ cells·mL^-1^ over a 60-72h period. Viruses isolated from warm waters took 17% longer than those from cold water to do so [38], suggesting that suboptimal hosts reduced the rate of infection as shown in cyanophages [68]. In contrast, infection dynamics of our isolates often resulted in host density of infected cultures growing to a steady state, but at a lower cell density than uninfected cells. (Supplementary Video 1), irrespective of cluster, population assignment or evidence of genes associated with temperate lifestyles. Out of 117 viruses isolated in this study, only 16 infections reduced host abundance below their inoculum density of 10^6^ cells·mL^-1^. In 53 infections, densities of infected cells increased to within an order of magnitude of uninfected cells (Supplementary Fig. 13), but demonstrated clear evidence of viral infection in cytograms, TEMs and subsequent recovery of viral genomes in selected samples.

Similar patterns of infection were recently reported in the extremely abundant bacteriophage ΦCrAss001 found in the human gut, where infection of exponentially growing cells of *B. intestinalis* 919/174 did not result in complete culture lysis, but caused a delay in stationary phase onset time and final density, despite lacking genes associated with lysogeny. As with our study, the authors observed that this only occurred in liquid culture, and isolation of the virus required numerous rounds of enrichment. They postulated that the virus may cause a successful infection in only a subset of host cells, with the remainder exhibiting alternative interactions such as pseudolysogeny or dormancy [32]. The prevalence of similar infection dynamics in the phages isolated in this study offer two intriguing possibilities: (i) Many of the viruses isolated in this study are either not fully lytic, but fall somewhere on the continuum of persistence [69]. This could be controlled by genes currently lacking a known function, with lysogeny (and associated superinfection immunity) favoured at high cell density, which would support the Piggyback-the-Winner hypothesis [70, 71]; (ii) the steady-state of host and virus densities observed here are an indicator of host phenotypic bistability in these streamlined heterotrophic taxa. Viral propagation occurring in only a subset of cells could explain the requirement of multiple rounds of enrichment before sufficient viral load is reached to be able to observe lytic infection on the host population. Either strategy, or a combination of both, would provide an ecological advantage of long-term stable coexistence between viruses and hosts and offer an explanation of the paradox of stable high abundances of both predator and prey across global oceans [12, 38, 72, 73]. Infection in a subset of the population could also explain the low lytic activity observed in pelagiphages *in situ*, despite high host densities [74]; and the small dynamic range and decoupled abundances of SAR11 and virioplankton in the Sargasso Sea [75]. Limited lysis of subpopulations of hosts such as SAR11 and OM43 that specialize in the uptake of labile carbon enriched through viral predation [76, 77] could facilitate efficient intra-population carbon recycling and explain the limited influence of SAR11 and associated viral abundances on carbon export to the deep ocean [5]. We propose the moniker the ‘Soylent Green Hypothesis’ for this mechanism, after the 1973 cult film in which the dead are recycled into food for the living. Further investigation leveraging our new virus-host model will provide greater insight into viral influence on ocean carbon biogeochemistry.

In conclusion, our method coupled Dilution-to-Extinction cultivation of hosts and associated viruses, resulting in the isolation of three new strains of OM43; a Western English Channel variant of a warm water ecotype of SAR11; the first known methylophages for OM43; the first siphovirus infecting SAR11, as well as eleven other viruses infecting this important marine heterotrophic clade and >100 more isolates to be sequenced and explored. The described method represents an efficient and costeffective strategy to isolate novel virus-host systems for experimental evaluation of co-evolutionary dynamics of important fastidious taxa from marine and other biomes. Coupling these methods to existing advances in host cultivation requires minimal additional effort and will provide valuable representative genomes to improve success rates of assigning putative hosts to metagenomically-derived viral contigs. Broader representation of model systems beyond cyanophages and viruses of copiotrophic, *r*-strategist hosts will reduce bias in developing methods to delineate viral population boundaries [78, 79], increasing the accuracy with which we derive ecological meaning from viral metagenomic data. We therefore hope that this method will enable viruses to be included in the current resurgence of cultivation efforts to better understand the biology and ecology of phages, and the influence of the world’s smallest predators on global biogeochemistry.

## Methods Summary

A complete description of the materials and methods is provided in the Supplementary Information. Four bacterial strains (*Methylophilales sp*. C6P1, D12P1 and H5P1; *Pelagibacter sp*. H2P3α) were isolated from Western English Channel station L4 seawater samples using Dilution-to-Extinction methods [29]. All four bacteria and two additional SAR11 strains *Pelagibacter bermudensis* HTCC7211 and *Pelagibacter ubique* HTCC1062 were used as bait to isolate phages from six monthly Western English Channel L4 seawater samples (50°15.00N; 4°13.00W). Briefly, water samples were concentrated for viruses using tangential flow filtration (100k Da Hydrosart membrane) and used as viral inoculum (10% v/v) in exponentially growing cultures of host bacteria in artificial seawater medium [80] in 96-well Teflon plates (Radleys, UK). Cells of the resulting lysate were filtered out (0.1 μm PVDF syringe filters) and the filtrate was used as viral inoculum in another round of isolation. This process was repeated until viral infection could be detected by flow cytometry - comparing cytograms and maximum density of infected cultures against uninfected cultures. Phages were purified by dilution-to-extinction methods (detailed protocol available here: dx.doi.org/10.17504/protocols.io.c36yrd). Phage genomes were sequenced using Illumina 2×150 PE sequencing, assembled and manually annotated as described in [81]. Phylogenetic classification of phages was performed on concatenated shared genes using a combination of Bayesian inference trees, maximum likelihood trees and shared-gene likelihood analyses, depending on the availability of appropriate taxa. ICTV-recognised genera based on shared gene content were assigned with VConTACT2 [19]. The relative abundance of novel phages in the Western English Channel [12] and Global Ocean viromes [11] was estimated (RPKM) by competitive read recruitment of five million randomly subsampled reads against pelagiphage and methylophage genomes from this study and others [38, 52, 64, 82].

## Data availability

All reads can be found in the SRA database under BioProject number PRJNA625644 as BioSamples SAMN14604128 - SAMN14604140. Annotated phage genomes are deposited as GenBank submissions under accession numbers MT375519 - MT375531. Sequences used for phylogenetic analysis are deposited under https://github.com/ViralPirates/viral-dte.

## Acknowledgements

We would like to thank Christian Hacker and the Bioimaging Centre of the University of Exeter for performing the TE microscopy and imaging. We would also like to thank the crew of the *R/V Plymouth Quest* and our collaborators at PML for collecting water samples, and the driver Magic for delivering water samples from Plymouth to Exeter. Genome sequencing was provided by MicrobesNG (http://www.microbesng.uk) which is supported by the BBSRC (grant number BB/L024209/1). This project used equipment funded by the Wellcome Trust Institutional Strategic Support Fund (WT097835MF), Wellcome Trust Multi-User Equipment Award (WT101650MA) and BBSRC LOLA award (BB/K003240/1). Bioinformatic analyses were conducted using the high-performance computing, ISCA, provided by the University of Exeter.

## Supplementary Information

### Supporting Methods

#### Water sampling

A total of 20 L of seawater was collected in rosette-mounted Niskin bottles at a depth of 5m from the Western Channel Observatory (WCO; http://www.westernchannelobservatory.org.uk/) coastal station ‘L4’ (50°15.00N; 4°13.00W) on the following dates: 2018-09-24, 2018-10-17, 2018-11-05, 2019-02-11, 2019-03-11, 2019-04-01, 2019-07-22 (for details see Supplementary Table 2). Seawater was transferred immediately to a clean 2 L acid-washed polycarbonate (PC) Nalgene bottles (Thermo Fisher Scientific, Waltham, USA) and placed in a cooler box (Igloo, Katy, USA) at ambient temperature. Upon return to shore, water was transported to the University of Exeter for immediate processing (Two hours maximum duration from collection to processing).

#### Isolation of SAR11 strain H2P3a and OM43 strains

1 L seawater was collected as described above in September 2017, filtered through a 142 mm Merck Millipore pore-size 0.2 μm PC filter and autoclaved in 2 L Duran bottles (DWK Life Sciences GmbH, Mainz, Germany) with the lid tightly shut to maintain carbon chemistry [1]. Upon cooling, nutrients were added to make natural seawater medium (NSW): 1 mM NH_4_Cl, 10 μM K_2_HPO_4_, 1 μM FeCl_3_, 25 μM glycine, 25 μM methionine, 100 μM pyruvate [2]. 1 mL of medium was placed into each 2 mL well in three sterile 96-well acid washed Teflon plates (Radleys, UK). Surface water from station L4 was filtered through a 142 mm Merck Millipore 0.2 μm PC filter to remove larger plankton and bacteria and enrich for smaller bacteria. A 200 μL aliquot was stained with SybrGreen and then quantified on a C6 Accuri flow cytometer. The remaining retained fraction was diluted in NSW to a density of ~ 1 cell·μL^-1^. 1 μL of diluted inoculum was added to each well, with eight wells left blank as negative controls and eight wells with undiluted low-nucleic acid community fraction as positive controls. Plates were covered and incubated in the dark at 15 °C for three months and checked monthly for positive growth by flow cytometry as described above. Positive wells (>10^6^ cells·mL^-1^) were transferred into 50 mL cultures in NSW medium and passaged three times once they achieved >10^6^ cells·mL^-1^). To verify culture purity and to taxonomically classify new isolates, the 16S rRNA gene was amplified from 10 μL of exponentially growing culture, heated to 95°C for 10 minutes, followed by PCR (25 cycles, 30 s @ 98 °C; 45 s @ 59 °C; 45 s @ 72 °C; final extension of 90 s @ 72 °C) using commercially available 27F and 1492R primers by Eurofins Genomics. Amplicon DNA was purified for Sanger sequencing (Eurofins) using a QIAquick PCR purification kit (Qiagen) following the manufacturer’s standard protocol. 16S rRNA sequences were compared to known bacterial sequences in the SILVA SSU database with SINA [3]. Sequences were aligned using T-coffee [4], curated using Gblocks [5] and maximum likelihood trees for both alignments were created using PhyML with standard settings and 500 bootstraps [6]. The trees were visualised using FigTree (v1.4.4, available https://github.com/rambaut/figtree).

#### Host cultivation

SAR11 strains HTCC1062 and HTCC7211 were kindly provided by the Giovannoni lab (Oregon State University, USA) and were kept in continuous culture in 50 mL polycarbonate flasks at 15 °C on artificial seawater medium ASM1 [7] throughout the duration of the study. Following isolation, cultures of OM43 and H2P3α were maintained on ASM1; medium for OM43 was further amended with 100 mM MeOH and 32 μL of commercially available MEM amino acid solution per 1.6 L of medium (Sigma, M5550), to improve cellular yields and reduce occurrences of chaining cells (observed through EM, Fig. 5, and a phenotype of alanine starvation in SAR11 [2]), respectively.

#### Viral concentration and inoculation

Monthly samples of 2 L of surface water were collected from station L4 as described above and sequentially filtered through a 142 mm Whatman GF/D filter (2.7 μm pore size) and a 142 mm 0.2 μm pore polycarbonate filter (Merck Millipore) using a peristaltic pump, to remove larger sized bacterioplankton. The filtrate was concentrated to 50 mL (40-fold concentration) using a 50R VivaFlow tangential flow filtration unit (Sartorius Lab instruments, Goettingen, Germany). The concentrate was filtered through a 0.1 μm pore PVDF membrane syringe-filter to remove any residual small cells and used as inoculum for viral isolation. 1 mL of exponentially growing bacterial host in ASM1 medium (amended with nutrients for OM43 strains as described for host cultivation above) was placed into each 2 mL well of a 96-well, acid washed Teflon plate. Eight wells were used as blank medium controls (no cell controls). 100 μL of viral inoculum was added to each well in the 96-well plate, with eight wells amended with 10% (v/v) medium instead of viral inoculum to serve as no-virus controls. The plates were incubated for ~2 weeks until no-virus controls reached maximum cell density (~2 weeks for strains HTCC1062, HTCCC7211 and H2P3α; about 1 week for OM43 strains H5P1, C6P1, D12P1). Growth was monitored at the end of the incubation period by flow cytometry; incubation periods were determined by the average growth times and data of the strains used for infections. Cytograms of virus-amended wells and no-virus controls were compared to identify infections as described below. A detailed protocol of the viral isolation process is available on www.protocols.io (DOI: dx.doi.org/10.17504/protocols.io.c36yrd). Once the no-virus controls reached maximum cell density, inoculated wells that did not show signs of infection were pooled (according to host type), filtered through a 0.1 μm pore PVDF membrane syringe (Durapore) filter to remove the cells and then used as fresh inoculum in a new 96-well plate of target host. This was repeated up to three times to amplify host-specific viral particles to a density whereby signs of viral infectivity could be observed on the flow cytometer.

#### Identification of positive infection via flow cytometry

We observed positive infections in cells via flow cytometry by comparing cytograms of wells inoculated with viruses against no-virus controls. Cytograms of wells containing infected cultures were identifiable by (1) a population shift of up to a 10-fold increase in green fluorescence compared to uninfected cultures; (2) an increase in the ‘noise’ fraction (presumably from cellular debris following lysis as well as viral particles) over time (Supplementary Video 1); and (3) a reduced maximum cell density (ranging from 10 to 1000-fold depending on initial cell density and viral inoculum concentration) in infected cultures compared to no-virus controls (Fig. 1B). The presence of viral-like particles in wells where a shift in green fluorescence, increased noise, and reduced maximum cell density were observed, were verified by TEM and reinfection of new cultures in fresh medium.

#### Viral purification

Following identification of a positive infection within a well, the contents of the well were filtered through a 0.1 μm pore-size PVDF syringe filter (Durapore) and used as inoculum in three rounds of viral purification. Briefly, a 96-well Teflon plate was inoculated with bacterial hosts as described previously. A 10-fold serial dilution series (from 10°-10^-10^) of the viral inoculum in ASM1 was added to a single row of the plate at 10 % v/v, with one well per row as no-virus controls (total of eight no-virus controls/plate). Plates were incubated at 18 °C for ~2 weeks until the no-virus controls reached maximum cell density and then screened for signs of viral infection using flow cytometry as described above. For each dilution series, the well amended with the lowest number of viruses that showed positive signs of infection was identified, and used as the inoculum in another round of viral purification [8]. Using this format, we were able to purify eight to twelve viral isolates simultaneously per plate.

#### Viral isolation costs and handling time

For each sample (processed environmental water sample), all steps of the initial viral isolation process (counting one to two plates for a total of three times) required ~6 hours run time on a flow cytometer. For the subsequent three rounds of purification, one plate was required for each host-sample combination (another ~6 hours of cytometer run time in total for one host-sample combination). Initial isolation of viruses from environmental water samples and three rounds of purification took ~10 weeks of incubation time in total. Between all steps, our protocol required ~7 hours of handling time per sample. Following three rounds of viral purification, generating sufficiently high viral titres to extract enough DNA for sequencing (approximately two weeks incubation time, and roughly four hours handling time over two days) was the rate-limiting step, and so required subselection of available viruses for sequencing. Future advances in DNA extraction efficiency, reducing DNA input requirements for sequencing and/or automation of viral DNA extraction will enable all isolated viruses to be sequenced. We estimate the cost of isolating a single virus is ~£20 for cultivation, flow cytometry and DNA extraction consumables, and ~£50-100 for genome sequencing at 30-fold coverage required for successful assembly, giving a total cost of £70-120 per sequenced viral isolate, not including the costs for person-hours. Thus, our protocol provides a high-throughput and scalable approach to viral isolation.

#### Transmission Electron Microscopy of viral isolates

For ultrastructural analysis, virus particles were transferred onto pioloform-coated electron microscopy (EM) copper grids (Agar Scientific, Standsted, UK) by floating the grids on droplets of virus-containing suspension for 3 min. Following a series of four washes on droplets of deionized water, the bound virus particles were contrasted with 2 % (w/v) uranyl acetate in 2 % (w/v) methyl cellulose (mixed 1:9) on ice for 8 min and the grids then air-dried on a wire loop after carefully removing excess staining solution with a filter. Dried grids were inspected with a JEOL JEM 1400 transmission electron microscope operated at 120 kV and images taken with a digital camera (ES 1000W CCD, Gatan, Abingdon, UK).

#### Host ranges of viral isolates

To test the infectivity of phage isolates against different hosts, 2 mL 96-well Teflon plates were prepared with 1 mL of exponentially growing bacterial host (HTCC1062, HTCC7211, H2P3α) in ASM1 medium. As medium control, ASM1 without any bacteria was added to eight wells. 100 μL of phage inoculum was added to eight wells per host type. As no-virus control, for each host further eight wells were left without any viruses added to them. The plate was incubated for ~2 weeks and analysed for signs of infection using flow cytometry as described above. A phage was classified as unable to infect a certain host if none of the eight amended replicates showed signs of infection.

#### DNA preparation and sequencing of viral isolates

50 mL ASM1 (amended with 100 mM MeOH and amino acid solution for OM43 as described above) in 250 mL acid-washed, polycarbonate flasks were inoculated with host cultures to 10^6^ cells·mL^-1^. The cultures were infected with 10% v/v viruses in ASM1 medium and incubated until after host cell lysis was detected using flow cytometry. The cultures were transferred to 50 mL Falcon tubes and the cellular fraction was removed by centrifugation (GSA rotor, Thermo Scientific 75007588) at 8,500 rpm/10,015 x g for 120 minutes. The supernatant was subsequently filtered through pore-size 0.1 μm PVDF syringe filter membranes to remove any remaining smaller cellular debris. Phages in the filtrate were precipitated using a modified version of an established PEG8000/NaCl DNA isolation method [9]. Briefly, the filtered phage lysate was transferred into 50 mL Falcon tubes with pre-weighted 5 g PEG8000 and 3.3 g NaCl, shaken until both dissolved and incubated on ice overnight. The phage particles were then pelleted by centrifugation at 8,500 rpm/10,015 x g for 90 min at 4 °C. Supernatant was discarded and phage particles resuspended by rinsing Falcon tubes twice with 1 mL SM buffer (100 mM NaCl, 8 mM MgSO4·7H2O, 50 mM Tris-Cl). DNA was extracted using the Wizard DNA Clean-Up system (Promega) following manufacturer’s instructions, but instead of TE we used pre-heated 60 °C nuclease free water to elute DNA. DNA was sequenced by MicrobesNG (Birmingham, UK) with Illumina paired-end HiSeq 2500. Default settings were used for all programs listed below, unless specified otherwise. All reads were trimmed, quality-controlled and error-corrected using bbmap and tadpole [10]. Contigs were assembled using SPAdes v3.13 [11] and evaluated with QUAST [12]. The trimmed reads were mapped back against the contigs for scaffolding using bowtie2 [13], BamM (alignment 0.9, identity 0.95) (available at https://github.com/minillinim/BamM) and samtools [14]. Viral contigs were confirmed with VirSorter (categories 1 or 2, >15kbp) and only accepted if the representative contig showed mean depth coverage an order of magnitude greater than the next highest recruiting contig (to filter out any cellular DNA carryover). Gene calls returned by VirSorter were imported into DNA Master for manual curation [15]. Additional gene calls were made using GenMark [16], GenMarkS [17], GenMarkS2 [18], GenMark.hmm [19], GenMark.heuristic [20], Glimmer v.3.02 [21], and Prodigal v.2.6.3 [22]. All gene calls were listed and compared using a scoring system which evaluates gene length, gene overlap and coding potential of ORFs [15]. ORFs were then annotated using BLASTp against the NCBI’s non-redundant protein sequences [23], Phmmer [24] against UniProts UniProtKB and uniprotrefprot [25], Swissprot [26] as well as InterProScan [27] and Pfam [28]. Virally-encoded tRNAs were identified with tRNAScan-SE v2.0 using settings `‘tRNAscan-SE −qQ --detail −o# −m# −f# −l# −c tRNAscan-SE.conf −s# −B’` [29] to identify putative tRNA genes. Matching tRNA genes (e.g. viral and host tRNA-Leu) were then aligned using BLASTn [30]. All viral genomes were organised into viral clusters (ICTV-recognised genera) based on shared gene content with VConTACT2 [31], and into populations using a boundary cutoff of 95% ANI over 85% contig length [32].

#### Evaluation of gene-sharing networks with hypergeometric testing

For the hypergeometric testing we gathered all annotated genomes from this study, other known pelagiphage isolates [33, 34] and we arbitrarily picked one representative contig from each cluster of complete phage genomes that were retrieved from a metagenomic Mediterranean deep chlorophyll maximum sample, that the authors speculated to be infective towards 1A SAR11s and included them in the phylogenetic analysis [35]. The contigs were imported into anvio v6.1 [36] where open reading frames were identified internally using Prodigal v2.6.3. The contigs were then used to run a pangenome analysis using the `--use-ncbi-blast` option. The binning function was then used to export the protein clusters, from which a matrix of the sum of shared proteins between all phages was created. The hypergeom function in the sciPy.stats python package was used to calculate the probability of sharing a protein between phages, based on which we created an average correlation cluster map.

#### Viral phylogenetic analysis

First we identified additional contigs to include in the phylogenetic analysis. The Global Ocean Virome (GOV2) [37] and a virome from the Western English Channel [38] were screened for contigs that belonged to the same population (95% ANI over 80% length using ClusterGenomes.pl - https://github.com/simroux/ClusterGenomes) or the same genus (clustered by VConTACT2 [39] into the same viral cluster, using default parameters) as isolate genomes. Additionally, all available genomes from Pelagibacter phage isolates were included, as well as selected fosmid derived contigs from Mediterranean metagenomes and 26 sequences from metagenomic mining that were identified as putative Pelagimyophages [33–35, 40, 41]. All sequences were subjected to gene calling using Prodigal v2.6.3. (default parameters) and screened for TerL genes by aligning all proteins against known TerL genes in annotated genomes (belonging to clusters A, B and D, as well as siphovirus *Kolga*) from this study using Protein BLAST (default parameter) [22, 30]. The TerL gene of Bordetella phage LK3 was added as an outgroup. The TerL genes were aligned within the Phylogeny.fr online service [42] opting for MUSCLE alignment [43] and Gblocks curation [5] with default settings. A maximum-likelihood tree was calculated with PhyML [44, 45], visualised using FigTree (v.1.4.4, available at http://tree.bio.ed.ac.uk/software/fiqtree/) and edited in Inkscape (www.inkscape.org) for aesthetics (font adjustments and addition of GOV2 metadata).

We only identified a single additional contig from metagenomes using the approach described above that was similar to OM43 phage *Venkman*. Only one other known phage for a related host (LD28 phage P19250A [46]) was available. These three contigs as well as all 14,883 viral contigs publicly available in January 2020 on http://millardlab.org/ were gene called using Prodigal v2.6.3. [22]. All called genes were compared to vFams [47] using hmmsearch [48] and sorted by E-value for best hits. The best matching gene sequences that were identified as similar to shared vFams found in *Venkman* and P19250A were important to the Phylogeny.fr online service [42]. Sequences of were aligned individually using T-coffee [4], curated using Gblocks [5] and concatenated manually. Maximum likelihood trees for both alignments were created using PhyML with standard settings and 500 bootstraps [6, 44]. The trees were visualised with FigTree (v.1.4.4) and edited in Inkscape (www.inkscape.org) for aesthetics. All sequences used for phylogenetic analysis are available at https://github.com/ViralPirates/viral-dte.

#### Calculation of Average Nucleotide Identity

All-vs-all average nucleotide identity (ANI) for our new isolates and existing pelagiphages using nucmer v.4.1 with the --nooptimize flag, followed by show-coords to convert the delta file into tabular format. ANI was calculated by taking the mean of all matching regions between two isolates.

#### Viral metagenomic recruitment

For assessing relative abundances of phage contigs in global datasets, a virome from the Western English Channel and all samples of the Global Ocean Virome dataset (GOV2) were used for recruitment [37, 38]. Metagenomic reads were downloaded from the European Nucleotide Archive ERR2625613 and subsampled to 5 million reads using bbmap’s reformat.sh command. A bowtie2 [13] index of dereplicated contigs was created from 25 known pelagiphage genomes from isolates [33, 34, 41], LD28 phage P19250A [46] and from this study six viral population representative genomes isolated on SAR11 hosts and one Methylophilales phage *Venkman* isolated on OM43 host. Reads were mapped against the genomes for each metagenome sample using bowtie2 using these commands: `bowtie2 --seed 42 --non-deterministic`. To calculate coverage and Reads Per Kilobase of contig per Million reads (RPKM) we used coverm [reference], with the following commands: `coverm contig -- bam-files *.bam --min-read-percent-identity 0.9 --methods rpkm --min-covered-fraction 0.4`. To minimize false positive detection rates [32], contigs that did not meet the 40% minimum genome coverage in a given sample were given an RPKM value of 0.

#### Evaluation of isolation efficiency as a function of matching host and virus geography

In total, 105 attempts were made for isolating viruses for SAR11 across the three host strains (HTCC7211: 35, HTCC1062: 40; H2P3α: 30). For each strain, the number of successful attempts were recorded in a vector, *v_s_*, of length *n* where *n* is the number of attempts made to isolate a virus, such that *v_i_*=1 if a successful attempt was made and *v_i_*=0 if an unsuccessful attempt was made. To test whether observed rates of success for H2P3 were significantly elevated compared to those for HTCC1062 and HTCC7211, the observed rates were evaluated against the null hypothesis that host strain did not make a difference to isolation success. Briefly, for each pair (H2P3 vs HTCC1062; H2P3α vs HTCC7211), the two vectors *v_s1_* and *v_s2_* were concatenated, and then randomly subsampled 999 times with replacement to a size of the smallest value of *n* between the vectors. The number of successes in each subsample were counted and the distribution of values across the 999 bootstraps was recorded. *P*-values were calculated by evaluating how many times within the 999 bootstraps a value as extreme as the observed value had been recorded.

#### 16S rRNA high-throughput amplicon sequencing and analysis

Genomic DNA was extracted using Qiagen DNeasy PowerWater Kit (REF 14900-50-NF) from biomass retained on 0.2 μm PC filters following the manufacturer’s protocol with minor modifications to increase the yield. Step 7 was modified from a 5 minute to 10 minute vortex bead beat. Step 21 was changed to have a 2 minute incubation with EB warmed to 55C. Amplification of the hypervariable V4 region of the 16S rRNA gene was performed using the 515fB (5’-GTGYCAGCMGCCGCGGTAA-3’) and 806rB (5’-GGACTACNVGGGTWTCTAAT-3’) primers. Libraries for each reaction were done attaching dual indices and Illumina adapters with the NexTera XT Index Kit (Illumina Inc.) Purified libraries were pooled and sequenced in a MiSeq platform. Primer sequences from paired-end fastq files were trimmed using CutAdapt [49]. Trimmed sequences were quality filtered, dereplicated and merged with DADA2 R package (version 1.8 [50]). Taxonomic assignment was performed using the “assignTaxonomy” command within dada2 pipeline and silva_nr_v132 database [51] as training set. SAR11 ASVs were extracted for further characterization using Phyloassigner v6.166 [52] and oligotyping [53]. SAR11 phylogenetic database SAR11_Phy_DB used in this study is available at https://www.github.com/lbolanos32/NAAMES_2020. Relative contribution barplots were done in R using ggplot2 [54] and edited in inkscape (www.inkscape.org) for aesthetics.

