## Supplementary Information for "Efficient Dilution-to-Extinction isolation of novel virus-host model systems for fastidious heterotrophic bacteria"

38. Warwick-Dugdale J, Solonenko N, Moore K, Chittick L, Gregory AC, Allen MJ, et al. Long-read

metagenomics reveals cryptic and abundant marine viruses. *bioRxiv* . 2018. , 345041

48. Eddy S. HMMER: profile HMMs for protein sequence analysis. 1998.

49. Martin M. Cutadapt removes adapter sequences from high-throughput sequencing reads. *EMBnet.journal* 2011; **17**: 10–12.

50. Callahan BJ, McMurdie PJ, Rosen MJ, Han AW, Johnson AJA, Holmes SP. DADA2: High-resolution sample inference from Illumina amplicon data. *Nat Methods* 2016; **13**: 581–583.

| | C6P1 | D12P1 | H5P1 | HTCC2181 | KB13 | HIMB624 | MBRSH7 |
|---|---|---|---|---|---|---|---|
| **C6P1** | | 98.24 | 98.31 | 96.17 | 95.00 | 94.10 | 95.06 |
| **D12P1** | 98.24 | | 98.83 | 96.62 | 95.39 | 94.50 | 95.46 |
| **H5P1** | 98.31 | 98.83 | | 97.79 | 96.55 | 95.80 | 96.62 |
| **HTCC2181** | 96.17 | 96.62 | 97.79 | | 96.49 | 95.40 | 96.55 |
| **KB13** | 95.00 | 95.39 | 96.55 | 96.49 | | 99.70 | 99.93 |
| **HIMB624** | 94.10 | 94.50 | 95.80 | 95.40 | 99.70 | | 99.60 |
| **MBRSH7** | 95.06 | 95.46 | 96.62 | 96.55 | 99.93 | 99.60 | |

| Collection Time | Collection Date | Temperature (° | Fluorescen | Depth (m) | Density | Salinity (ps | CStarTr0 | Par | Oxy |
|---|---|---|---|---|---|---|---|---|---|
| 10:21:14 | Sept 24 2018 | 15.546 | 1.6909 | 5 | 1025.87 | 34.9962 | 85.2961 | 0.4093 | 239.931 |
| 09:20:37 | Oct 17 2018 | 14.8181 | 0.9033 | 5 | 1025.709 | 34.5798 | 69.7736 | 2.1833 | 236.302 |
| 10:27:16 | Nov 05 2018 | 14.1766 | 0.9475 | 5 | 1026.269 | 35.126 | 88.7715 | 0.6285 | 240.449 |
| 10:17:04 | Feb 11 2019 | 9.3449 | 0.6315 | 5 | 1026.494 | 34.2436 | 79.7166 | 1.6396 | 269 |
| 10:13:55 | Mar 11 2019 | 10.001 | 0.5425 | 5 | 1026.989 | 35.0152 | 89.5834 | 0.6723 | 267.411 |
| 09:21:13 | Apr 01 2019 | 10.394 | 0.5298 | 5 | 1026.958 | 35.0672 | 89.4219 | 0.4996 | 270.531 |
| 09:20:27 | Jul 22 2019 | 17.3155 | 0.5612 | 5 | 1025.634 | 35.2245 | 94.7492 | 0.1291 | 237.998 |
| 09:09:43 | Aug 12 2019 | 16.9301 | 0.5437 | 5 | 1025.753 | 35.2596 | 92.0995 | 0.2526 | 242.956 |

| Water Sample | Targeted host | Rounds of Enrichment |
|---|---|---|
| September-1 | 1062 | 2 |
| October-1 | 7211 | 3 |
| October-1 | 1062 | 3 |
| October-1 | H2P3alpha | 3 |
| October-1 | H5P1 | 3 |
| November | 7211 | 2 |
| November | 1062 | 2 |
| November | H2P3alpha | 2 |
| November | H5P1 | 2 |
| February | 7211 | 1 |
| February | 1062 | 1 |
| February | H2P3alpha | 1 |
| February | H5P1 | 2 |
| March | 7211 | 1 |
| March | 1062 | 1 |
| March | H2P3alpha | 1 |
| March | H5P1 | 2 |
| April | 7211 | 1 |
| April | 1062 | 1 |
| April | H2P3alpha | 1 |
| April | H5P1 | 2 |
| July | D12P1 | 2 |

| | Aegir | Bylgja | Eistla | Eyrgjafa | Gjalp | Greip | Himinglaeva | Hroenn | Jormungand | Kolga | Ran | Unn | HTVC010P | HTVC011P | HTVC019P | HTVC021P | HTVC022P | HTVC025P | HTVC031P | HTVC105P | HTVC109P | HTVC119P | HTVC120P | HTVC121P | HTVC200P | HTVC201P | uvMED-CGF-C97-MedDCM-OCT-S27-C11 | uvMED-CGR-C23A-MedDCM-OCT-S43-C49 | uvMED-CGR-C23B-MedDCM-OCT-S39-C49 | uvMED-CGR-C24-MedDCM-OCT-S38-C43 | uvMED-CGR-C26B-MedDCM-OCT-S35-C55 | uvMED-CGR-C28-MedDCM-OCT-S23-C36 | uvMED-CGR-C43-MedDCM-OCT-S32-C95 | uvMED-GF-C93-MedDCM-OCT-S32-C16 | uvMED-GF-C93-MedDCM-OCT-S38-C9 | uvMED-GF-U-MedDCM-OCT-S28-C24 | uvMED-GF-U-MedDCM-OCT-S32-C31 |
|---|---|---|---|---|---|---|---|---|---|---|---|---|---|---|---|---|---|---|---|---|---|---|---|---|---|---|---|---|---|---|---|---|---|---|---|---|---|
| Aegir | | | | | | | | | | 99.7 | | | | | | | | | | | | | | | | | | | | | | | | | | | |
| Bylgja | | | 80.7 | | | | 100.0 | 100.0 | 82.3 | 91.6 | 82.3 | 100.0 | | | 82.4 | | 82.4 | | 81.9 | 86.7 | 78.4 | 84.9 | 90.4 | 88.3 | 77.5 | | 82.1 | | | | | | | 83.0 | | | |
| Eistla | | 80.7 | | 93.1 | 93.1 | 95.2 | 80.7 | 80.7 | 84.8 | | 84.8 | 80.7 | | 91.6 | 83.3 | 81.1 | 84.3 | 85.5 | 86.9 | 82.1 | 81.9 | 84.0 | 80.9 | 80.8 | | 83.1 | | | | | | | | 81.5 | 81.4 | | |
| Eyrgjafa | | | 93.2 | | 100.0 | | | | | | | | | 89.1 | 93.3 | | 92.0 | 82.0 | 89.2 | | | 85.7 | | | | 94.3 | | | | | | | | | | | 78.8 |
| Gjalp | | | 93.2 | 100.0 | | | | | | | | | | 89.1 | 93.3 | | 92.0 | 82.0 | 89.2 | | | 85.7 | | | | 94.3 | | | 100.0 | | | | | | | | 78.8 |
| Greip | | | 95.2 | | | | | | | | | | 81.7 | 90.6 | | | 85.3 | | 89.8 | | | | | | | 90.2 | | 86.8 | 87.3 | 81.5 | 85.0 | 80.3 | | | | | |
| Himinglaeva | | 100.0 | 80.7 | | | | | 100.0 | 82.3 | 91.6 | 82.3 | 100.0 | | | 82.4 | | 82.4 | | 81.9 | 86.7 | 78.4 | 84.9 | 90.4 | 88.3 | 77.5 | | 82.1 | | | | | | | 83.0 | | | |
| Hroenn | | 100.0 | 80.7 | | | | 100.0 | | 82.3 | 91.6 | 82.3 | 100.0 | | | 82.4 | | 82.4 | | 81.9 | 86.7 | 78.4 | 84.9 | 90.1 | 88.3 | 77.5 | | 82.1 | | | | | | | 83.0 | | | |
| Jormungand | | 82.3 | 84.8 | | | | 82.3 | 82.3 | | 93.0 | 100.0 | 82.3 | | | 90.0 | 83.8 | 90.6 | | 82.4 | 83.6 | 79.6 | 87.3 | 86.2 | 83.6 | | 87.2 | | | | | | | | 84.1 | 84.3 | | |
| Kolga | 99.7 | 91.6 | | | | | 91.6 | 91.6 | 93.0 | | 93.0 | 91.6 | | | | | | | | | | 90.9 | 92.0 | | | | | | | | | | | | | | |
| Ran | | 82.3 | 84.8 | | | | 82.3 | 82.3 | 100.0 | 93.0 | | 82.3 | | | 89.6 | 83.7 | 90.4 | | 82.4 | 83.6 | 79.6 | 87.3 | 86.2 | 83.6 | | 87.3 | | | | | | | | 84.1 | 84.1 | | |
| Unn | | 100.0 | 80.7 | | | | 100.0 | 100.0 | 82.3 | 91.6 | 82.3 | | | | 82.4 | | 82.4 | | 81.9 | 86.7 | 78.4 | 84.9 | 90.1 | 88.3 | 77.5 | | 82.1 | | | | | | | 83.0 | | | |
| HTVC010P | | | | | | 82.3 | | | | | | | | | 80.9 | | | | | | | | | | | | | 84.0 | 83.8 | 84.3 | 85.3 | 89.6 | | | | | |
| HTVC011P | | | 90.8 | 88.6 | 88.6 | 90.6 | | | | | | | | | 92.9 | | 92.7 | | 92.5 | | | 86.9 | | | 86.5 | 86.2 | | 89.9 | 91.1 | | | | | | | | |
| HTVC019P | | 81.3 | 84.7 | 92.3 | 92.3 | | 81.3 | 81.3 | 89.5 | | 89.4 | 81.3 | | 92.8 | | 84.5 | 89.8 | | 84.7 | 84.5 | 82.2 | | 79.5 | 84.4 | | 85.1 | 84.3 | | | | | | | 83.7 | 84.4 | | |
| HTVC021P | | | 82.4 | | | | | | 83.3 | | 83.1 | | | | 84.5 | | 84.0 | | 84.9 | 88.6 | 82.1 | 88.5 | | 89.4 | | 83.6 | | | | | | | | 89.9 | 89.8 | | |
| HTVC022P | | 81.7 | 83.3 | 92.7 | 92.7 | 84.4 | 81.7 | 81.7 | 89.7 | | 89.4 | 81.7 | | 92.7 | 91.7 | 83.7 | | | 85.0 | 83.4 | 79.5 | 85.1 | 81.3 | 83.5 | 84.9 | 87.8 | | | | | | | | 84.2 | 84.3 | | |
| HTVC025P | | | 84.4 | 81.9 | 81.9 | | | | | | | | | | | | | | 85.9 | | | | | | | | | | | | | | | | | 80.1 | |
| HTVC031P | | 80.7 | 85.7 | 87.8 | 87.8 | 88.4 | 80.7 | 80.7 | 84.1 | | 84.1 | 80.7 | | 90.6 | 84.8 | 84.5 | 85.2 | 86.1 | | 84.8 | 79.1 | | 80.8 | 83.1 | 80.5 | 82.9 | 80.0 | 88.8 | 88.8 | | | | | 82.6 | 82.6 | | 83.8 |
| HTVC105P | | | 82.5 | | | | | | 82.6 | | 82.4 | | | | 83.3 | 88.3 | 83.7 | | 84.8 | | 81.5 | | 86.2 | 91.6 | 86.5 | 82.4 | | | | | | | | 89.7 | 88.8 | | |
| HTVC109P | | | 80.6 | | | | | | | | | | | | | 81.5 | | | 80.9 | 81.3 | | | | 81.0 | | | | | | | | | | | | | |
| HTVC119P | | 87.5 | 83.1 | 89.2 | 89.2 | | 87.5 | 87.5 | 84.7 | 89.6 | 84.7 | 87.5 | | 86.6 | | 88.4 | 84.5 | | | 83.3 | | | 85.6 | 87.8 | 84.5 | 91.0 | | | | | | | | 85.2 | 85.8 | | 89.0 |
| HTVC120P | | 89.4 | 80.0 | | | | 89.4 | 89.1 | 88.7 | 90.6 | 88.7 | 89.1 | | | | 83.0 | 82.4 | | 81.1 | 86.4 | | 85.4 | | 81.5 | 81.8 | | 84.3 | | | | | | | | 87.6 | | |
| HTVC121P | | | 82.4 | | | | | | 83.7 | | 83.7 | | | | 83.2 | 89.0 | 84.2 | | 83.9 | 89.9 | 80.8 | | | | | 82.7 | | | | | | | | 88.4 | 90.2 | | |
| HTVC200P | | 81.2 | | | | | 81.2 | 81.2 | | | | 81.2 | | | | | 85.4 | | 81.6 | 94.6 | | 84.4 | 81.8 | | | | | 85.2 | 90.4 | | | | | | 86.6 | | 84.4 |
| HTVC201P | | | 83.0 | 92.8 | 92.8 | 88.8 | | | 86.3 | | 86.0 | | | 87.0 | 84.9 | 83.5 | 86.4 | | 83.4 | 82.7 | | 91.1 | 83.4 | 83.2 | | | | | | | | | | 83.4 | 84.0 | | |
| uvMED-CGF-C97-MedDCM-OCT-S27-C11 | | 82.5 | | | | | 82.5 | 82.5 | | | | 82.5 | | | 84.3 | | | | | | 86.0 | | 84.5 | | | | | | | | | | | | | | 88.6 |
| uvMED-CGR-C23A-MedDCM-OCT-S43-C49 | | | | | | 84.7 | | | | | | | 84.8 | 89.8 | | | | | 88.8 | | | | | | 84.7 | | | | 96.3 | 81.8 | | 84.8 | | | | 84.1 | |
| uvMED-CGR-C23B-MedDCM-OCT-S39-C49 | | | | | 99.0 | 85.8 | | | | | | | 83.4 | 91.1 | | | | | 88.8 | | | | | | | | | 96.3 | | 86.6 | | 84.5 | | | | 84.1 | |
| uvMED-CGR-C24-MedDCM-OCT-S38-C43 | | | | | 98.4 | 81.6 | | | | | | | 84.9 | | | | | | | | | | | | | | | 85.8 | 92.2 | | 84.1 | 86.8 | 84.7 | | | | |
| uvMED-CGR-C26B-MedDCM-OCT-S35-C55 | | | | | | 87.0 | | | | | | | 82.4 | | | | | | 88.0 | | | | | | | | | 88.3 | 88.3 | 84.1 | | 80.6 | | | | | |
| uvMED-CGR-C28-MedDCM-OCT-S23-C36 | | | | | | 83.9 | | | | | | | 90.0 | | | | | | | | | | | | | | | 84.6 | 84.3 | 87.8 | 81.3 | | 80.9 | | | | |
| uvMED-CGR-C43-MedDCM-OCT-S32-C95 | | | | | | | | | | | | | | | | | | | | | | | | | | | | | | 84.5 | | 84.7 | | | | | |
| uvMED-GF-C93-MedDCM-OCT-S32-C16 | | 81.6 | | | | | 81.6 | 81.6 | 83.6 | | 83.6 | 81.6 | | | 84.0 | 89.6 | 84.1 | | 82.8 | 89.6 | | 85.2 | 79.3 | 89.0 | | 83.5 | | | | | | | | | 94.3 | | |
| uvMED-GF-C93-MedDCM-OCT-S38-C9 | | | 84.6 | | | | | | 83.9 | | 83.7 | | | | 83.3 | 88.6 | 84.1 | | 82.6 | 89.0 | | 85.8 | 87.2 | 89.7 | 85.6 | 83.7 | | | 89.9 | | | | | 92.4 | | | |
| uvMED-GF-U-MedDCM-OCT-S32-C31 | | | | | | | | | | | | | | 89.0 | | | | | | | | 90.8 | | | 84.1 | | 89.7 | | | | | | | | | | |

| Exeter Viral Collectio | Isolate name | Viral Population Repr | viral genus (vContact | Hypergeometric clus | Other vContact2 clus | Reference | Genome | % completeness (che | Genome_length(bp) | %GC_content | Virus_Family | Host_strain | Host_clade |
|---|---|---|---|---|---|---|---|---|---|---|---|---|---|
| EXVC025P | Eistla | Eistla | A | Cluster A | Himinglaeva,Hroenn, | This Study, Zhao 201 | Circular | 95.37 | 39638 | 32.7 | Podoviridae | HTCC1062 | SAR11 1a |
| EXVC018P | Eyrgjafa | Eyrgjafa | A | Cluster B | Himinglaeva,Hroenn, | This Study, Zhao 201 | Circular | 100 | 38005 | 32.6 | Podoviridae | HTCC1062 | SAR11 1a |
| EXVC020P | Gjalp | Eyrgjafa | A | Cluster B | Himinglaeva,Hroenn, | This Study, Zhao 201 | Circular | 99.77 | 37857 | 32.5 | Podoviridae | HTCC1062 | SAR11 1a |
| EXVC021P | Greip | Greip | B | Cluster D | - | - | Circular | 101 | 34916 | 31.5 | Podoviridae | HTCC1062 | SAR11 1a |
| EXVC014P | Ran | Ran | A | Cluster A | Himinglaeva,Hroenn, | This Study, Zhao 201 | Circular | 99.62 | 41529 | 34.1 | Podoviridae | H2P3α | SAR11 1a |
| EXVC013S | Aegir | Kolga | C | Singleton | Aegir, Kolga | - | Fragment | 38.85 | 18297 | 31.1 | Siphoviridae | H2P3α | SAR11 1a |
| EXVC012P | Jörmungand | Ran | A | Cluster A | Himinglaeva,Hroenn, | This Study, Zhao 201 | Circular | 99.62 | 41529 | 34.1 | Podoviridae | H2P3α | SAR11 1a |
| EXVC016S | Kólga | Kolga | C | Singleton | Aegir, Kolga | - | Circular | 103.24 | 48659 | 30.5 | Siphoviridae | H2P3α | SAR11 1a |
| EXVC019P | Unn | Bylgja | A | Cluster A | Himinglaeva,Hroenn, | This Study, Zhao 201 | Circular | 98.1 | 41069 | 33.5 | Podoviridae | H2P3α | SAR11 1a |
| EXVC015P | Hroenn | Bylgja | A | Cluster A | Himinglaeva,Hroenn, | This Study, Zhao 201 | Circular | 98.13 | 41069 | 33.5 | Podoviridae | H2P3α | SAR11 1a |
| EXVC010P | Bylgja | Bylgja | A | Cluster A | Himinglaeva,Hroenn, | This Study, Zhao 201 | Circular | 98.1 | 41069 | 33.5 | Podoviridae | H2P3α | SAR11 1a |
| EXVC011P | Himinglæva | Bylgja | A | Cluster A | Himinglaeva,Hroenn, | This Study, Zhao 201 | Circular | 98.1 | 41069 | 33.5 | Podoviridae | H2P3α | SAR11 1a |
| EXVC028S | Venkman | Venkman | D | Singleton | P19250A | Moon 2017 | Linear | 103.86 | 38624 | 34.4 | Siphoviridae | H5P1 | OM43 |

| Exeter Viral Collectio | Enrichment rounds r | BioSample | BioProject | Accession_no | SRA | Collection Time | Collection_Date | Temperature (°C) | Fluorescence | Depth (m) |
|---|---|---|---|---|---|---|---|---|---|---|
| EXVC025P | 3 | SAMN14604129 | PRJNA625644 | MT375521 | SRR11559273 | 09:20:37 | Oct 17 2018 | 14.8181 | 0.9033 | 5 |
| EXVC018P | 3 | SAMN14604130 | PRJNA625644 | MT375523 | SRR11559269 | 09:20:37 | Oct 17 2018 | 14.8181 | 0.9033 | 5 |
| EXVC020P | 3 | SAMN14604131 | PRJNA625644 | MT375524 | SRR11559268 | 09:20:37 | Oct 17 2018 | 14.8181 | 0.9033 | 5 |
| EXVC021P | 3 | SAMN14604132 | PRJNA625644 | MT375525 | SRR11559267 | 09:20:37 | Oct 17 2018 | 14.8181 | 0.9033 | 5 |
| EXVC014P | 3 | SAMN14604133 | PRJNA625644 | MT375530 | SRR11559266 | 09:20:37 | Oct 17 2018 | 14.8181 | 0.9033 | 5 |
| EXVC013S | 3 | SAMN14604134 | PRJNA625644 | MT375519 | SRR11559265 | 09:20:37 | Oct 17 2018 | 14.8181 | 0.9033 | 5 |
| EXVC012P | 3 | SAMN14604135 | PRJNA625644 | MT375528 | SRR11559264 | 09:20:37 | Oct 17 2018 | 14.8181 | 0.9033 | 5 |
| EXVC016S | 3 | SAMN14604136 | PRJNA625644 | MT375529 | SRR11559263 | 09:20:37 | Oct 17 2018 | 14.8181 | 0.9033 | 5 |
| EXVC019P | 2 | SAMN14604137 | PRJNA625644 | MT375531 | SRR11559262 | 10:27:16 | Nov 05 2018 | 14.1766 | 0.9475 | 5 |
| EXVC015P | 2 | SAMN14604138 | PRJNA625644 | MT375527 | SRR11559272 | 10:27:16 | Nov 05 2018 | 14.1766 | 0.9475 | 5 |
| EXVC010P | 2 | SAMN14604139 | PRJNA625644 | MT375520 | SRR11559271 | 10:27:16 | Nov 05 2018 | 14.1766 | 0.9475 | 5 |
| EXVC011P | 2 | SAMN14604140 | PRJNA625644 | MT375526 | SRR11559270 | 10:27:16 | Nov 05 2018 | 14.1766 | 0.9475 | 5 |
| EXVC028S | 3 | SAMN14604128 | PRJNA625644 | MT375522 | SRR11559274 | 09:20:37 | Oct 17 2018 | 14.8181 | 0.9033 | 5 |

| Exeter Viral Collectio | Density | Salinity (psu) | CStarTr0 | Par | Oxy | Environment_broad | Environment_local | Latitude | Longitude |
|---|---|---|---|---|---|---|---|---|---|
| EXVC025P | 1025.7093 | 34.5798 | 69.7736 | 2.1833 | 236.302 | Marine | Coastal | 50◦15.00N | 4◦13.00W |
| EXVC018P | 1025.7093 | 34.5798 | 69.7736 | 2.1833 | 236.302 | Marine | Coastal | 50◦15.00N | 4◦13.00W |
| EXVC020P | 1025.7093 | 34.5798 | 69.7736 | 2.1833 | 236.302 | Marine | Coastal | 50◦15.00N | 4◦13.00W |
| EXVC021P | 1025.7093 | 34.5798 | 69.7736 | 2.1833 | 236.302 | Marine | Coastal | 50◦15.00N | 4◦13.00W |
| EXVC014P | 1025.7093 | 34.5798 | 69.7736 | 2.1833 | 236.302 | Marine | Coastal | 50◦15.00N | 4◦13.00W |
| EXVC013S | 1025.7093 | 34.5798 | 69.7736 | 2.1833 | 236.302 | Marine | Coastal | 50◦15.00N | 4◦13.00W |
| EXVC012P | 1025.7093 | 34.5798 | 69.7736 | 2.1833 | 236.302 | Marine | Coastal | 50◦15.00N | 4◦13.00W |
| EXVC016S | 1025.7093 | 34.5798 | 69.7736 | 2.1833 | 236.302 | Marine | Coastal | 50◦15.00N | 4◦13.00W |
| EXVC019P | 1026.2688 | 35.126 | 88.7715 | 0.6285 | 240.449 | Marine | Coastal | 50◦15.00N | 4◦13.00W |
| EXVC015P | 1026.2688 | 35.126 | 88.7715 | 0.6285 | 240.449 | Marine | Coastal | 50◦15.00N | 4◦13.00W |
| EXVC010P | 1026.2688 | 35.126 | 88.7715 | 0.6285 | 240.449 | Marine | Coastal | 50◦15.00N | 4◦13.00W |
| EXVC011P | 1026.2688 | 35.126 | 88.7715 | 0.6285 | 240.449 | Marine | Coastal | 50◦15.00N | 4◦13.00W |
| EXVC028S | 1025.7093 | 34.5798 | 69.7736 | 2.1833 | 236.302 | Marine | Coastal | 50◦15.00N | 4◦13.00W |

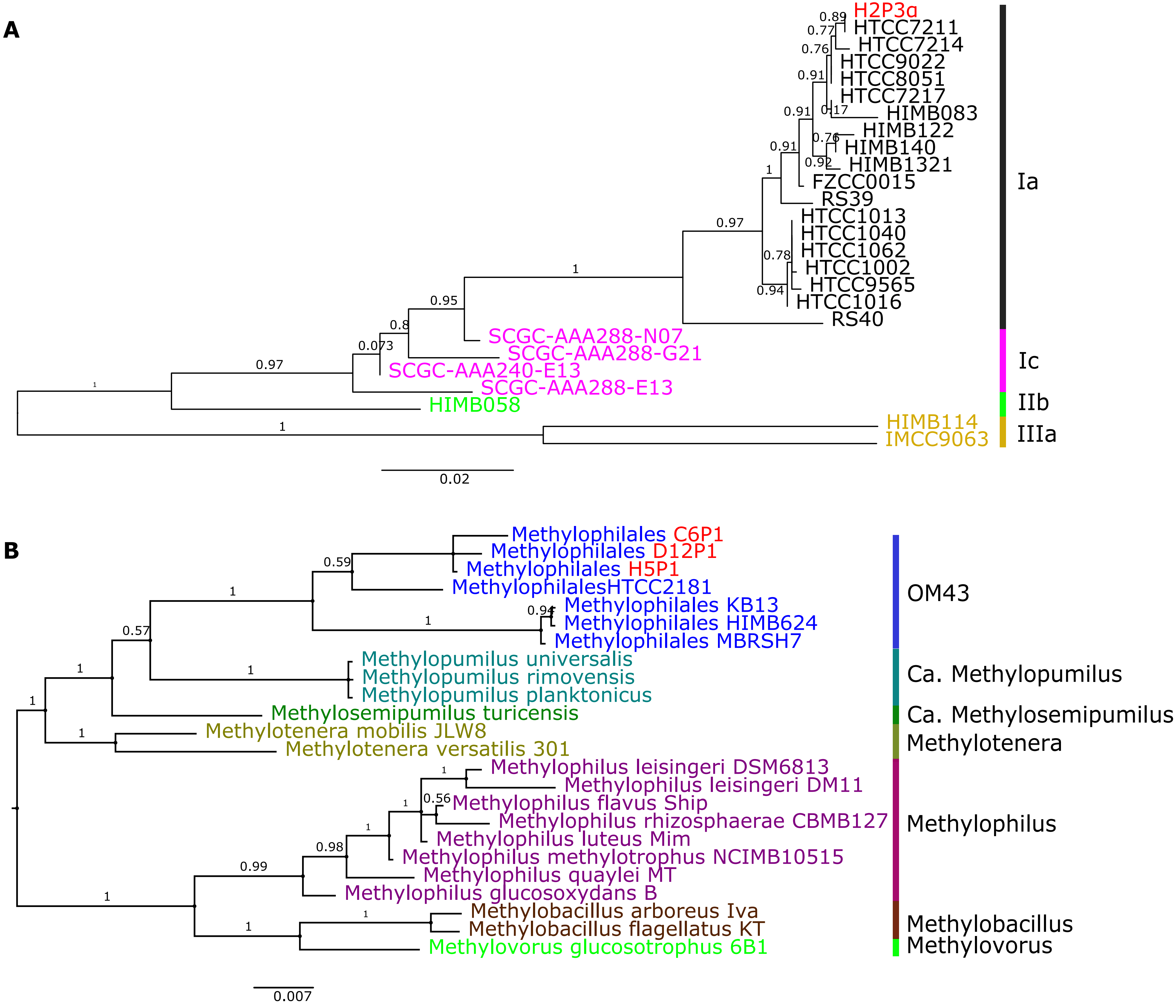
A
H2P3α
HTCC7211
HTCC7214
HTCC9022
HTCC8051
HTCC7217
HIMB083
HIMB122
HIMB140
HIMB1321
FZCC0015
RS39
HTCC1013
HTCC1040
HTCC1062
HTCC1002
HTCC9565
HTCC1016
RS40
SCGC-AAA288-N07
SCGC-AAA288-G21
SCGC-AAA240-E13
SCGC-AAA288-E13
HIMB058
HIMB114
IMCC9063
Ia
Ic
IIb
IIIa
0.02
B
Methylophilales C6P1
Methylophilales D12P1
Methylophilales H5P1
MethylophilalesHTCC2181
Methylophilales KB13
Methylophilales HIMB624
Methylophilales MBRSH7
Methylopumilus universalis
Methylopumilus rimovensis
Methylopumilus planktonicus
Methylosemipumilus turicensis
Methylotenera mobilis JLW8
Methylotenera versatilis 301
Methylophilus leisingeri DSM6813
Methylophilus leisingeri DM11
Methylophilus flavus Ship
Methylophilus rhizosphaerae CBMB127
Methylophilus luteus Mim
Methylophilus methylotrophus NCIMB10515
Methylophilus quaylei MT
Methylophilus glucosoxydans B
Methylobacillus arboreus Iva
Methylobacillus flagellatus KT
Methylovorus glucosotrophus 6B1
OM43
Ca. Methylopumilus
Ca. Methylosemipumilus
Methylotenera
Methylophilus
Methylobacillus
Methylovorus
0.007

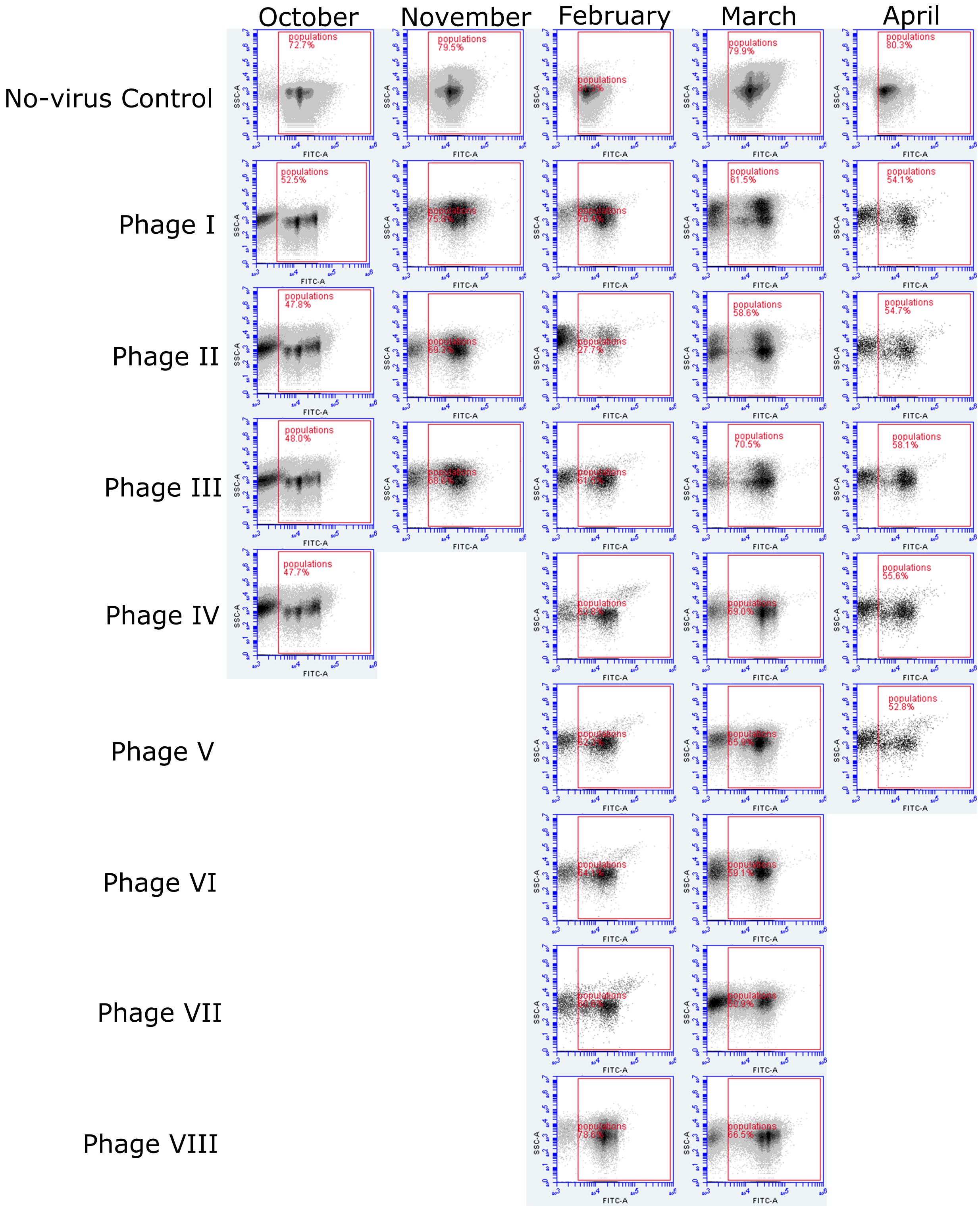

October
November
February
March
April
No-virus Control
Phage I
Phage II
Phage III
Phage IV
Phage V
Phage VI
Phage VII
Phage VIII
populations
SSC-A
FITC-A

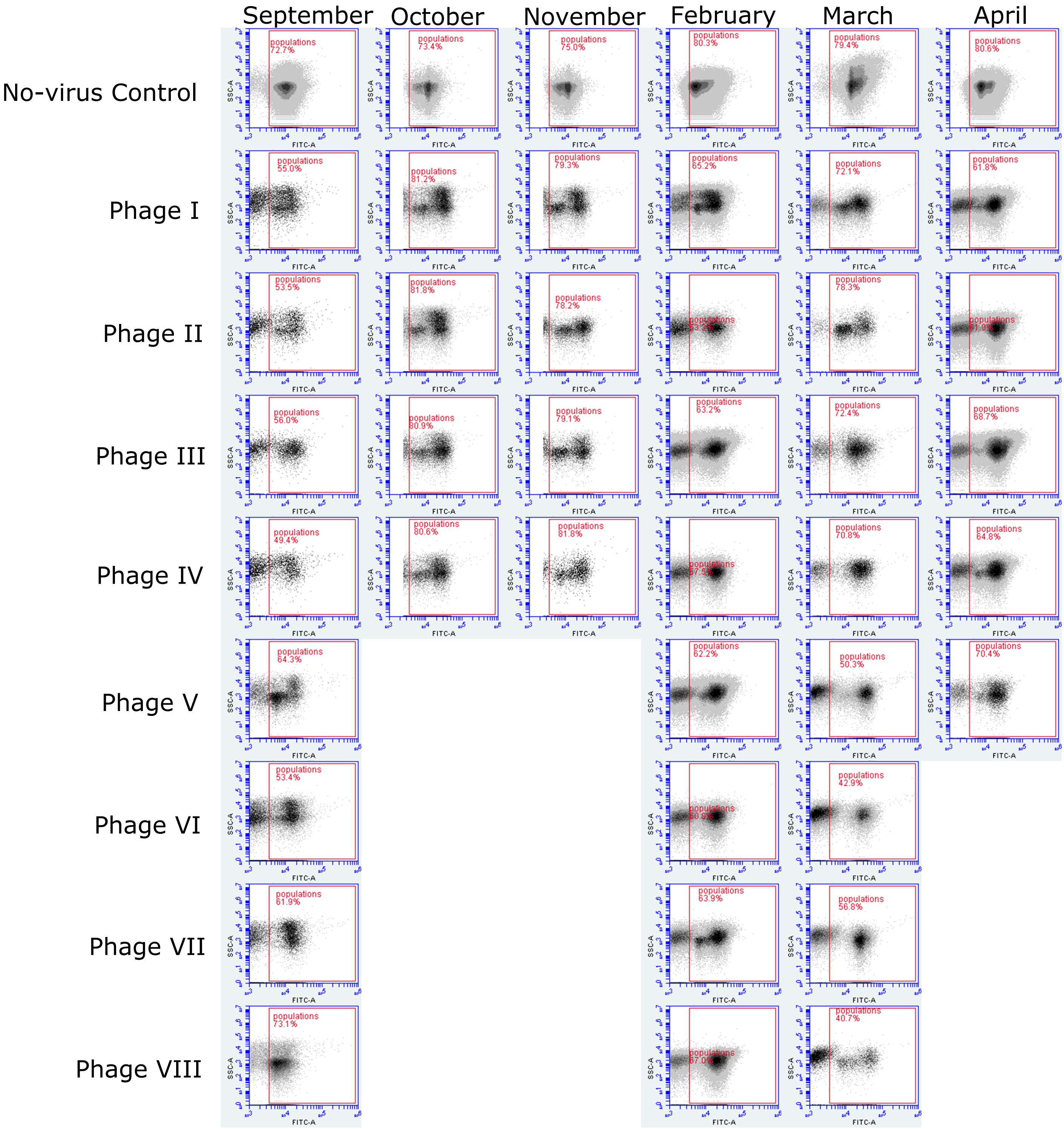
September
October
November
February
March
April
No-virus Control
Phage I
Phage II
Phage III
Phage IV
Phage V
Phage VI
Phage VII
Phage VIII
SSC-A
FITC-A
populations
72.7%
73.4%
75.0%
80.3%
79.4%
80.6%
55.0%
81.2%
79.3%
65.2%
72.1%
61.8%
53.5%
81.8%
78.2%
78.3%
56.0%
80.9%
79.1%
63.2%
72.4%
68.7%
49.4%
80.6%
81.8%
70.8%
64.8%
64.3%
62.2%
50.3%
70.4%
53.4%
42.9%
61.9%
63.9%
56.8%
73.1%
40.7%

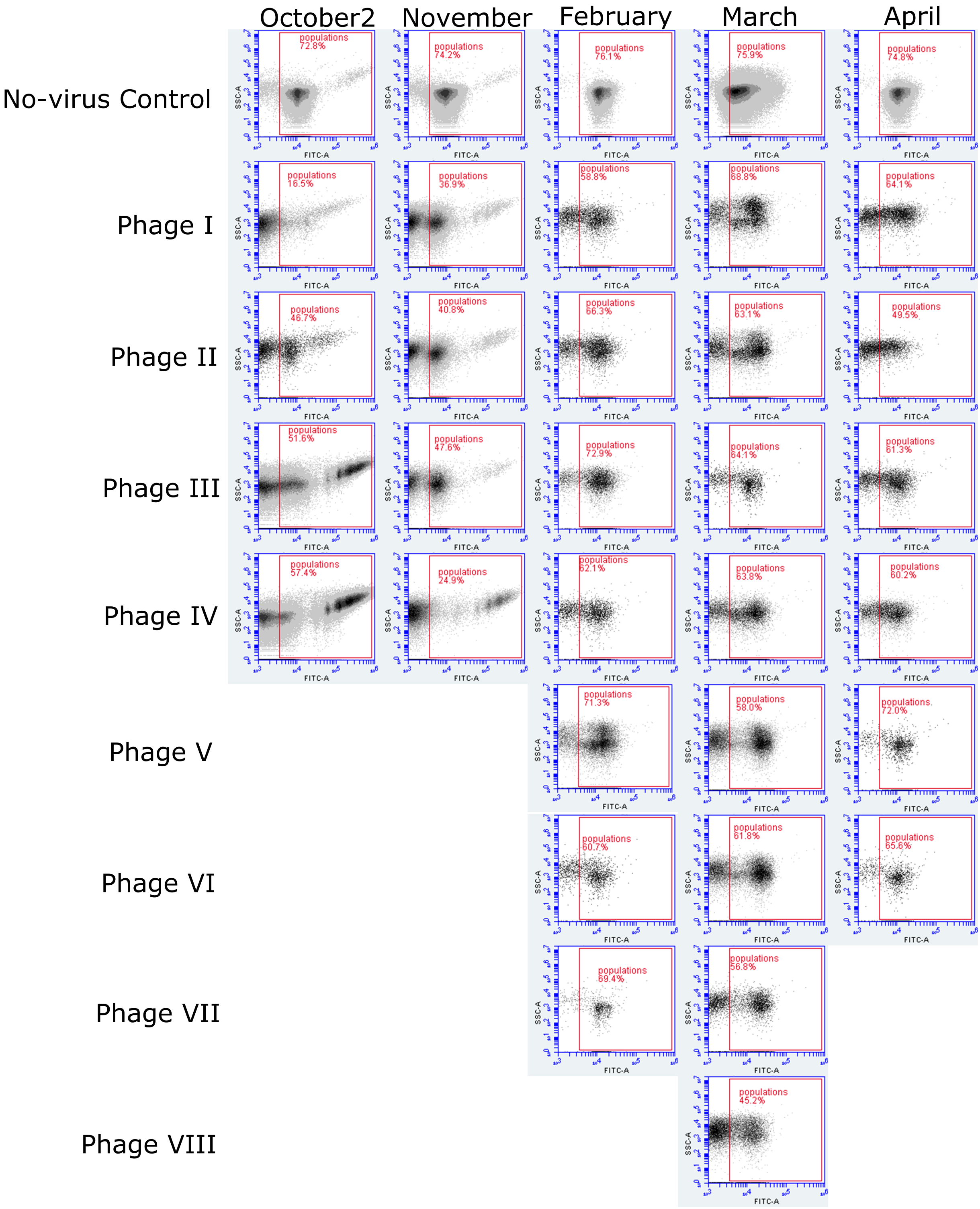

October2
November
February
March
April
No-virus Control
populations 72.8%
populations 74.2%
populations 76.1%
populations 75.9%
populations 74.8%
Phage I
populations 16.5%
populations 36.9%
populations 58.8%
populations 68.8%
populations 64.1%
Phage II
populations 46.7%
populations 40.8%
populations 66.3%
populations 63.1%
populations 49.5%
Phage III
populations 51.6%
populations 47.6%
populations 72.9%
populations 64.1%
populations 61.3%
Phage IV
populations 57.4%
populations 24.9%
populations 62.1%
populations 63.8%
populations 60.2%
Phage V
populations 71.3%
populations 58.0%
populations 72.0%
Phage VI
populations 60.7%
populations 61.8%
populations 65.6%
Phage VII
populations 69.4%
populations 56.8%
Phage VIII
populations 45.2%
SSC-A
FITC-A

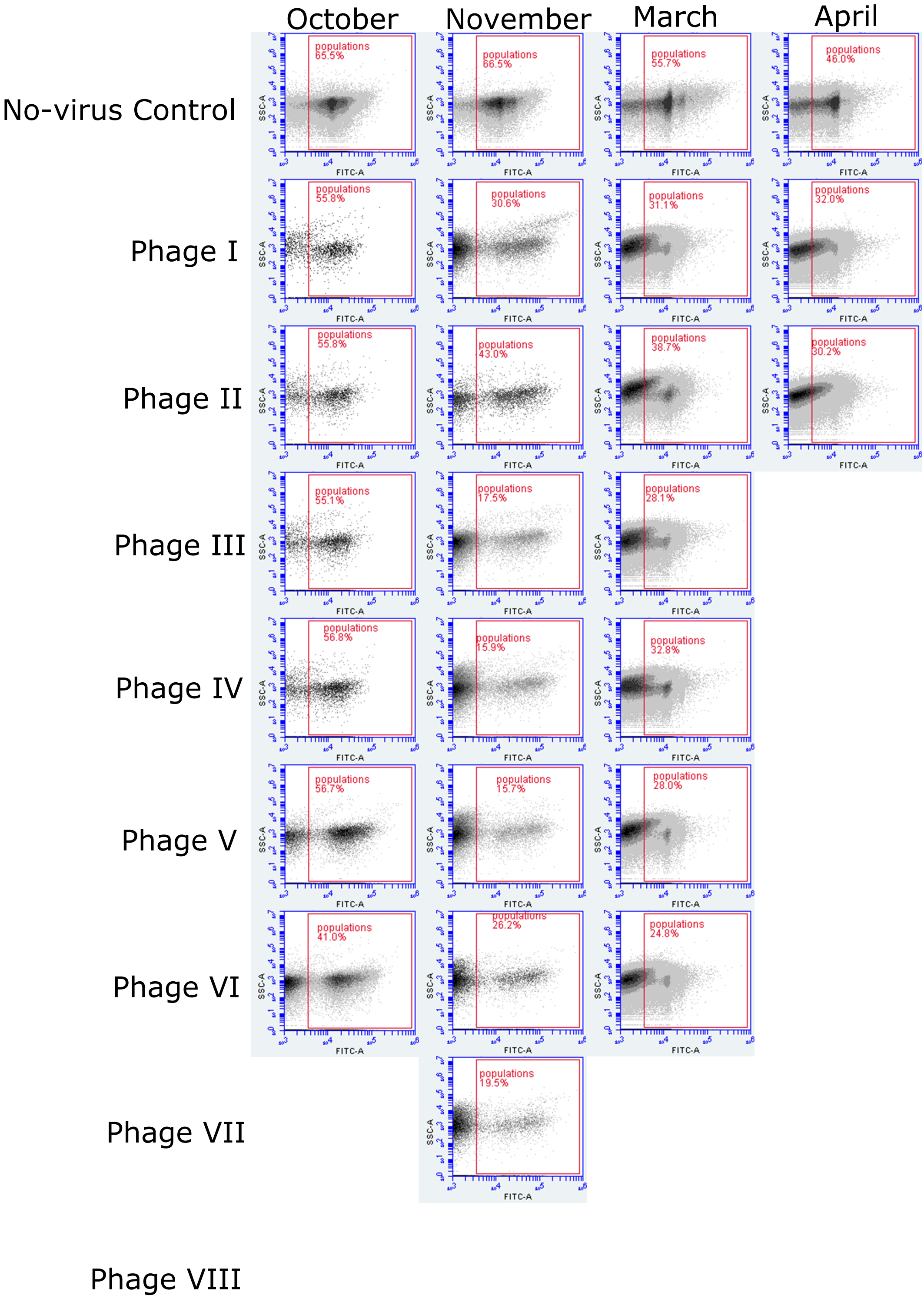

October
November
March
April
No-virus Control
Phage I
Phage II
Phage III
Phage IV
Phage V
Phage VI
Phage VII
Phage VIII
populations 65.5%
populations 66.5%
populations 55.7%
populations 46.0%
populations 55.8%
populations 30.6%
populations 31.1%
populations 32.0%
populations 55.8%
populations 43.0%
populations 38.7%
populations 30.2%
populations 55.1%
populations 17.5%
populations 28.1%
populations 56.8%
populations 15.9%
populations 32.8%
populations 56.7%
populations 15.7%
populations 28.0%
populations 41.0%
populations 26.2%
populations 24.8%
populations 19.5%
SSC-A
FITC-A

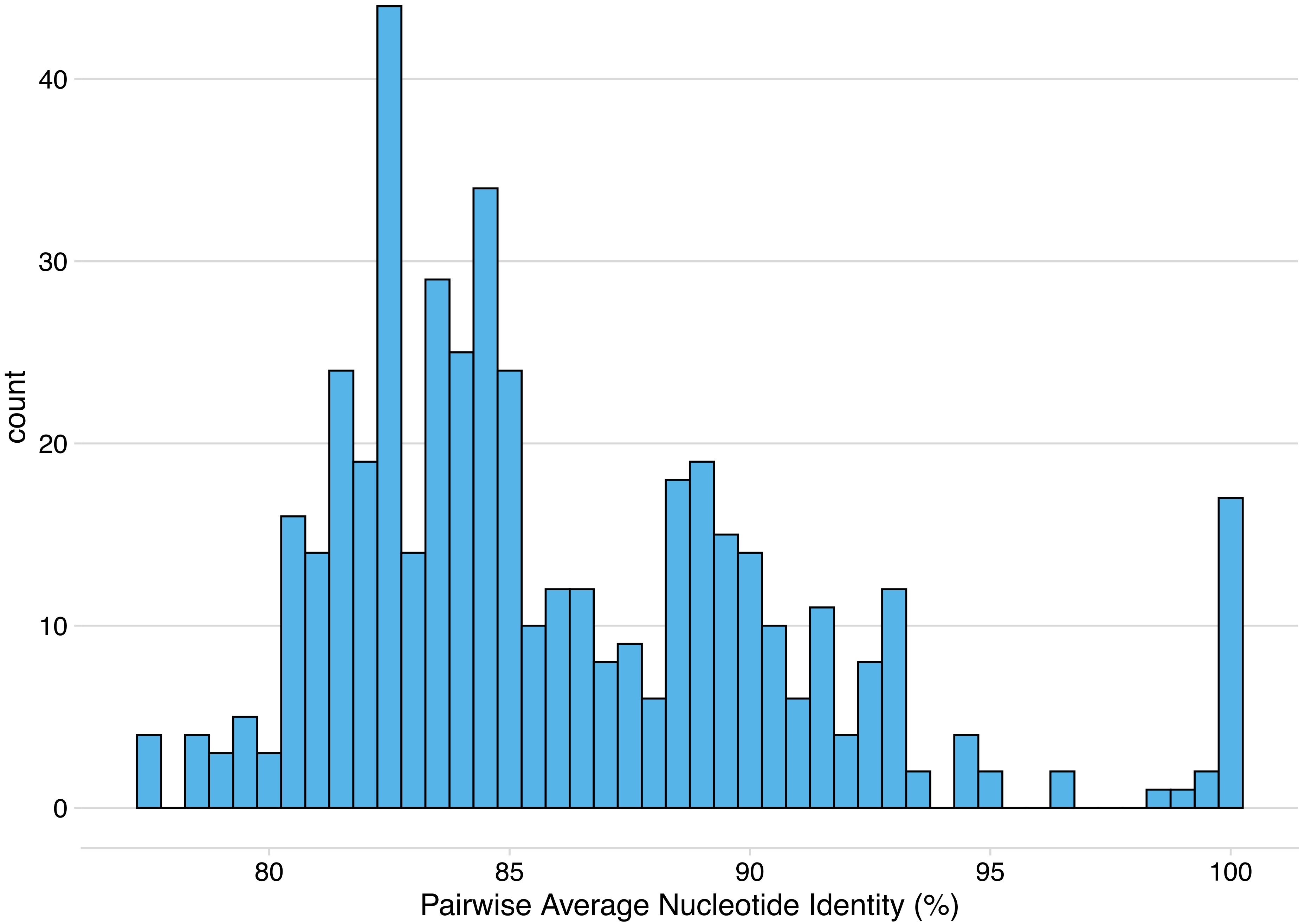
count
40
30
20
10
0
80
85
90
95
100
Pairwise Average Nucleotide Identity (%)

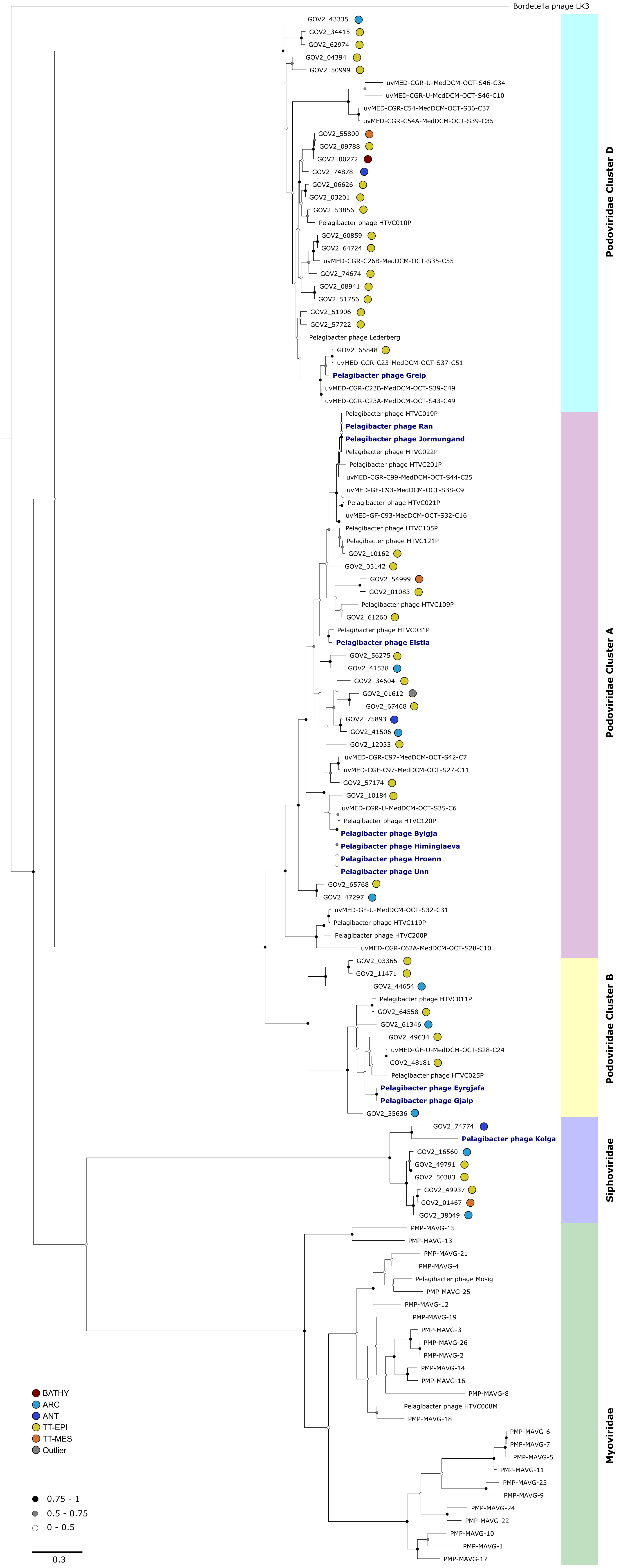

Bordetella phage LK3
GOV2_43335
GOV2_34415
GOV2_62974
GOV2_04394
GOV2_50999
uvMED-CGR-U-MedDCM-OCT-S46-C34
uvMED-CGR-U-MedDCM-OCT-S46-C10
uvMED-CGR-C54-MedDCM-OCT-S36-C37
uvMED-CGR-C54A-MedDCM-OCT-S39-C35
GOV2_55800
GOV2_09788
GOV2_00272
GOV2_74878
GOV2_06626
GOV2_03201
GOV2_53856
Pelagibacter phage HTVC010P
GOV2_60859
GOV2_64724
uvMED-CGR-C26B-MedDCM-OCT-S35-C55
GOV2_74674
GOV2_08941
GOV2_51756
GOV2_51906
GOV2_57722
Pelagibacter phage Lederberg
GOV2_65848
uvMED-CGR-C23-MedDCM-OCT-S37-C51
Pelagibacter phage Greip
uvMED-CGR-C23B-MedDCM-OCT-S39-C49
uvMED-CGR-C23A-MedDCM-OCT-S43-C49
Pelagibacter phage HTVC019P
Pelagibacter phage Ran
Pelagibacter phage Jormungand
Pelagibacter phage HTVC022P
Pelagibacter phage HTVC201P
uvMED-CGR-C99-MedDCM-OCT-S44-C25
uvMED-GF-C93-MedDCM-OCT-S38-C9
Pelagibacter phage HTVC021P
uvMED-GF-C93-MedDCM-OCT-S32-C16
Pelagibacter phage HTVC105P
Pelagibacter phage HTVC121P
GOV2_10162
GOV2_03142
GOV2_54999
GOV2_01083
Pelagibacter phage HTVC109P
GOV2_61260
Pelagibacter phage HTVC031P
Pelagibacter phage Eistla
GOV2_56275
GOV2_41538
GOV2_34604
GOV2_01612
GOV2_67468
GOV2_75893
GOV2_41506
GOV2_12033
uvMED-CGR-C97-MedDCM-OCT-S42-C7
uvMED-CGF-C97-MedDCM-OCT-S27-C11
GOV2_57174
GOV2_10184
uvMED-CGR-U-MedDCM-OCT-S35-C6
Pelagibacter phage HTVC120P
Pelagibacter phage Bylgja
Pelagibacter phage Himinglaeva
Pelagibacter phage Hroenn
Pelagibacter phage Unn
GOV2_65768
GOV2_47297
uvMED-GF-U-MedDCM-OCT-S32-C31
Pelagibacter phage HTVC119P
Pelagibacter phage HTVC200P
uvMED-CGR-C62A-MedDCM-OCT-S28-C10
GOV2_03365
GOV2_11471
GOV2_44654
Pelagibacter phage HTVC011P
GOV2_64558
GOV2_61346
GOV2_49634
uvMED-GF-U-MedDCM-OCT-S28-C24
GOV2_48181
Pelagibacter phage HTVC025P
Pelagibacter phage Eyrgjafa
Pelagibacter phage Gjalp
GOV2_35636
GOV2_74774
Pelagibacter phage Kolga
GOV2_16560
GOV2_49791
GOV2_50383
GOV2_49937
GOV2_01467
GOV2_38049
PMP-MAVG-15
PMP-MAVG-13
PMP-MAVG-21
PMP-MAVG-4
Pelagibacter phage Mosig
PMP-MAVG-25
PMP-MAVG-12
PMP-MAVG-19
PMP-MAVG-3
PMP-MAVG-26
PMP-MAVG-2
PMP-MAVG-14
PMP-MAVG-16
PMP-MAVG-8
Pelagibacter phage HTVC008M
PMP-MAVG-18
PMP-MAVG-6
PMP-MAVG-7
PMP-MAVG-5
PMP-MAVG-11
PMP-MAVG-23
PMP-MAVG-9
PMP-MAVG-24
PMP-MAVG-22
PMP-MAVG-10
PMP-MAVG-1
PMP-MAVG-17
Podoviridae Cluster D
Podoviridae Cluster A
Podoviridae Cluster B
Siphoviridae
Myoviridae
BATHY
ARC
ANT
TT-EPI
TT-MES
Outlier
0.75 - 1
0.5 - 0.75
0 - 0.5
0.3

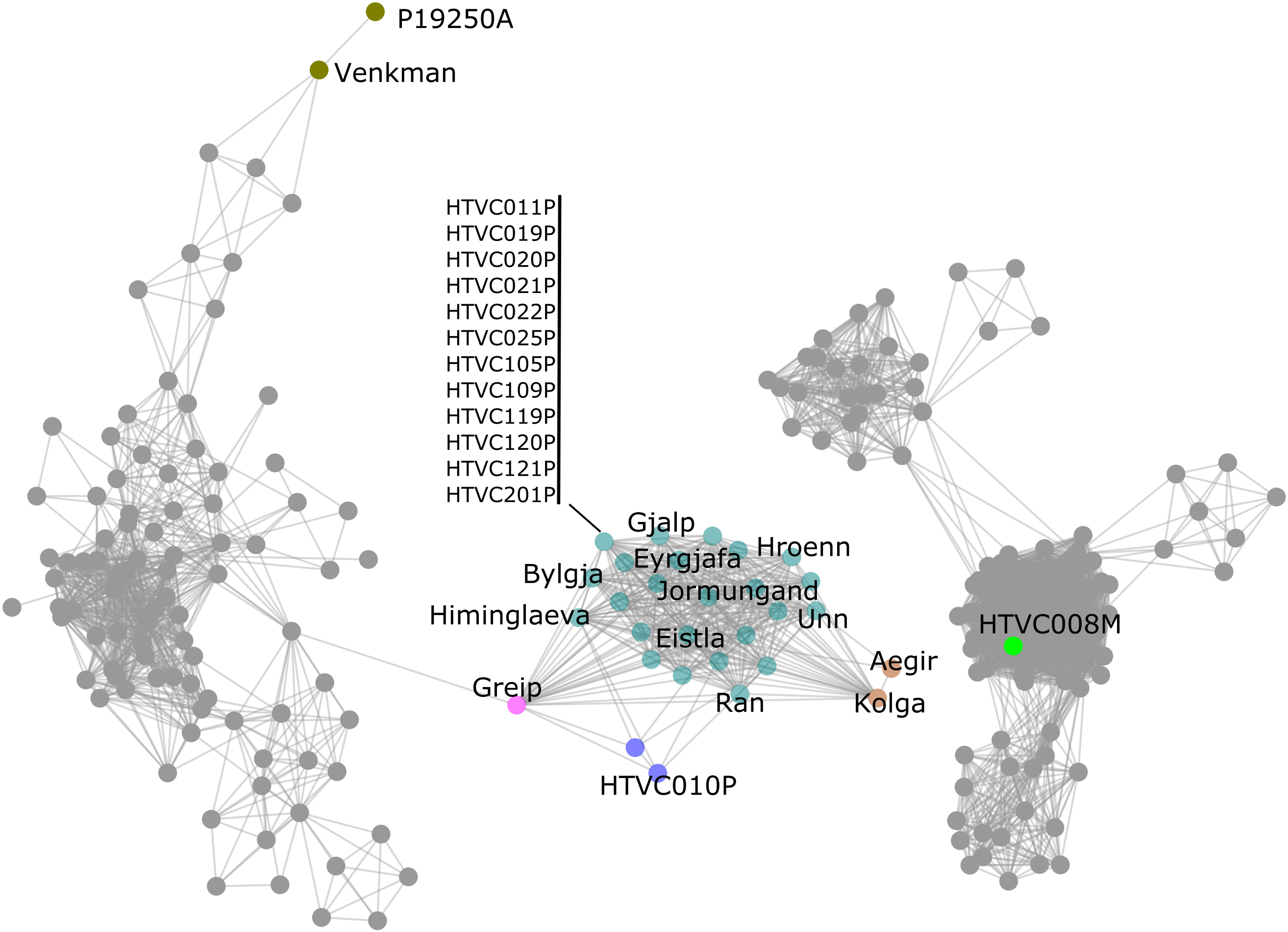
P19250A
Venkman
HTVC011P
HTVC019P
HTVC020P
HTVC021P
HTVC022P
HTVC025P
HTVC105P
HTVC109P
HTVC119P
HTVC120P
HTVC121P
HTVC201P
Gjalp
Hroenn
Eyrgjafa
Bylgja
Jormungand
Himinglaeva
Unn
Eistla
HTVC008M
Aegir
Greip
Ran
Kolga
HTVC010P

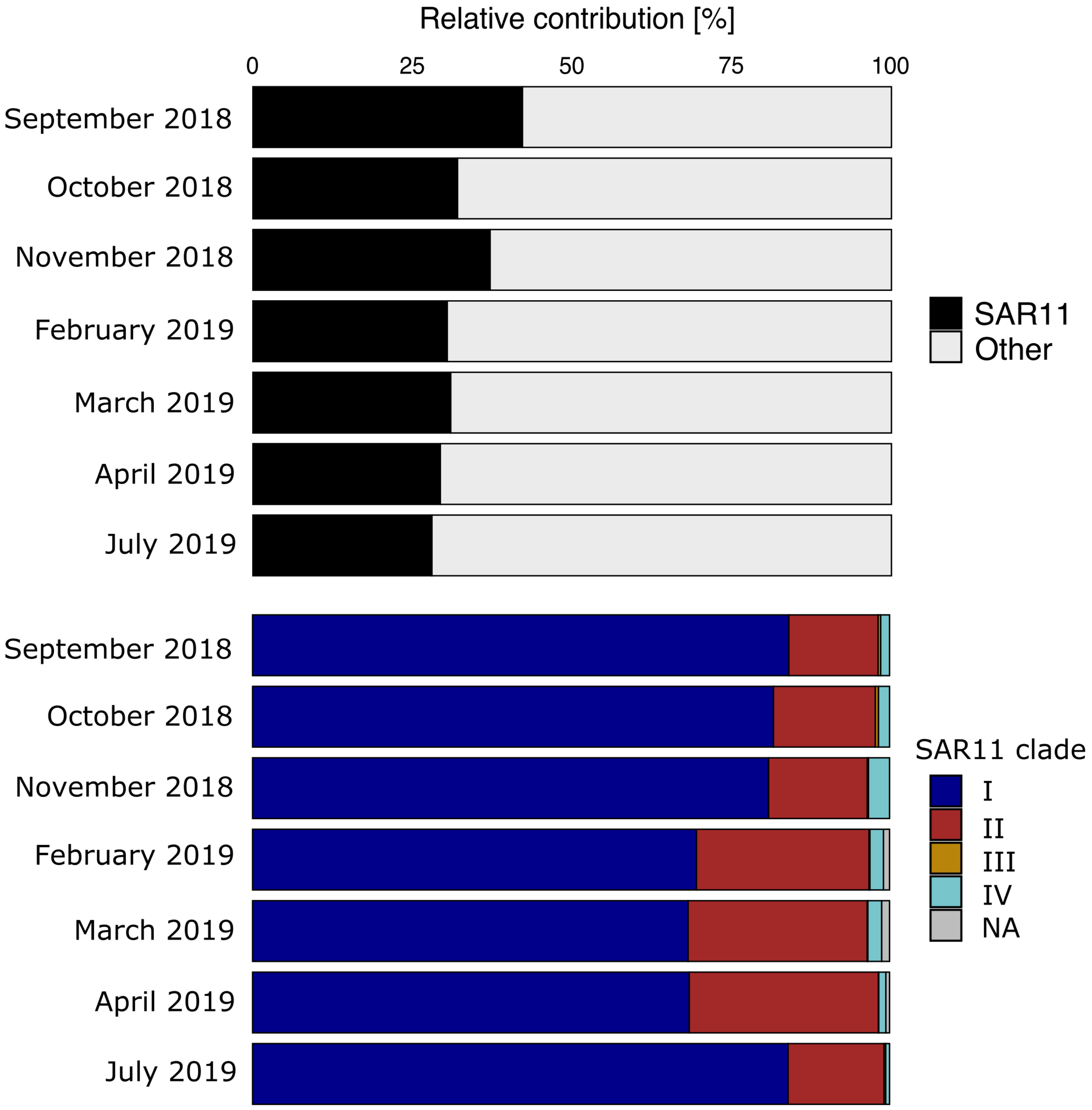

Relative contribution [%]
0
25
50
75
100
September 2018
October 2018
November 2018
February 2019
March 2019
April 2019
July 2019
SAR11
Other
SAR11 clade
I
II
III
IV
NA

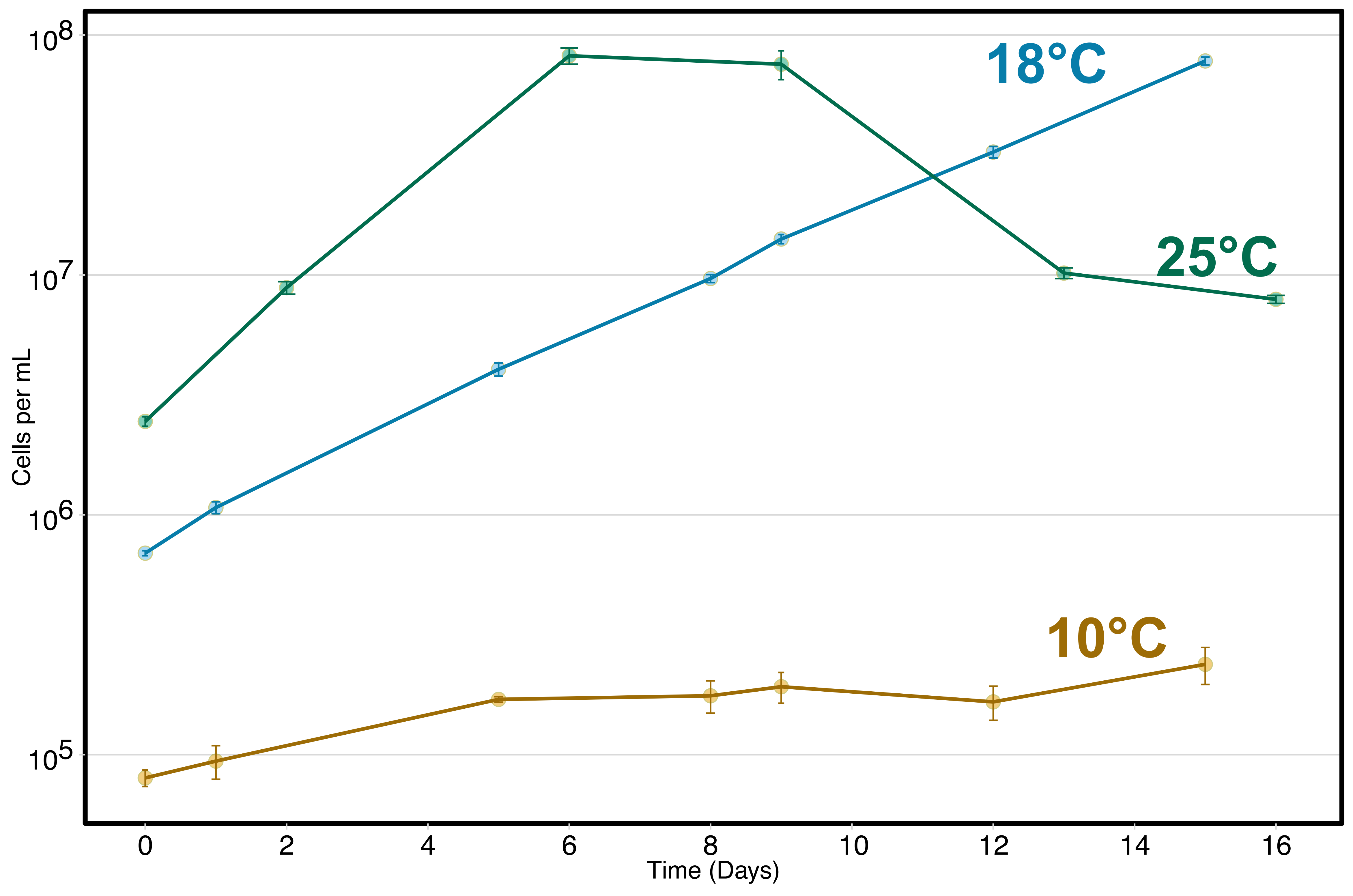

18°C
25°C
10°C
$10^8$
$10^7$
$10^6$
$10^5$
Cells per mL
0
2
4
6
8
10
12
14
16
Time (Days)

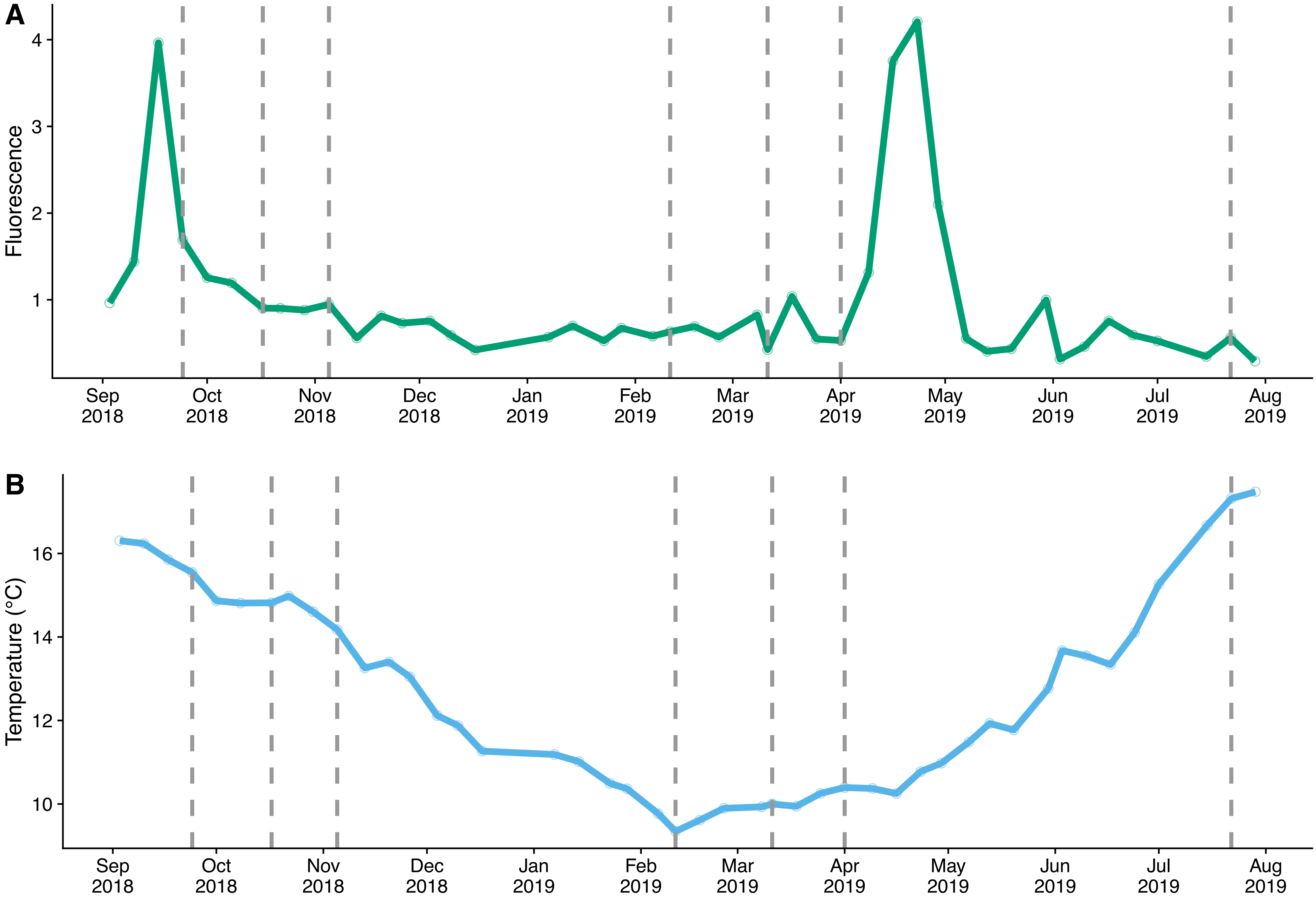
A
B
Fluorescence
1
2
3
4
Temperature (°C)
10
12
14
16
Sep 2018
Oct 2018
Nov 2018
Dec 2018
Jan 2019
Feb 2019
Mar 2019
Apr 2019
May 2019
Jun 2019
Jul 2019
Aug 2019

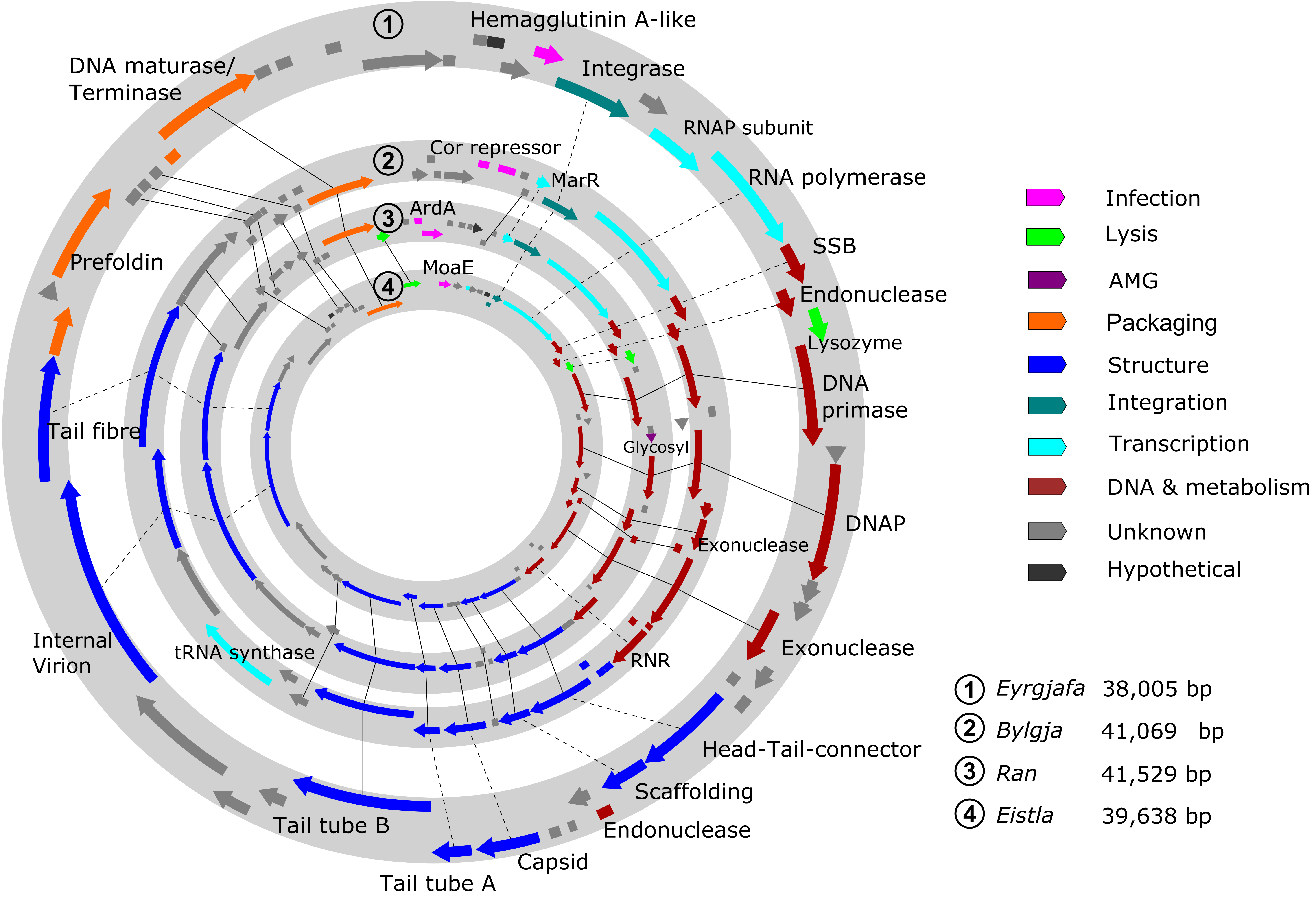

1
Hemagglutinin A-like
Integrase
RNAP subunit
RNA polymerase
SSB
Endonuclease
Lysozyme
DNA primase
DNAP
Exonuclease
Exonuclease
Head-Tail-connector
Scaffolding
Endonuclease
Capsid
Tail tube A
Tail tube B
Internal Virion
tRNA synthase
Tail fibre
Prefoldin
DNA maturase/ Terminase
2
Cor repressor
MarR
3
ArdA
4
MoaE
Glycosyl
RNR
Infection
Lysis
AMG
Packaging
Structure
Integration
Transcription
DNA & metabolism
Unknown
Hypothetical
1 Eyrgjafa 38,005 bp
2 Bylgja 41,069 bp
3 Ran 41,529 bp
4 Eistla 39,638 bp

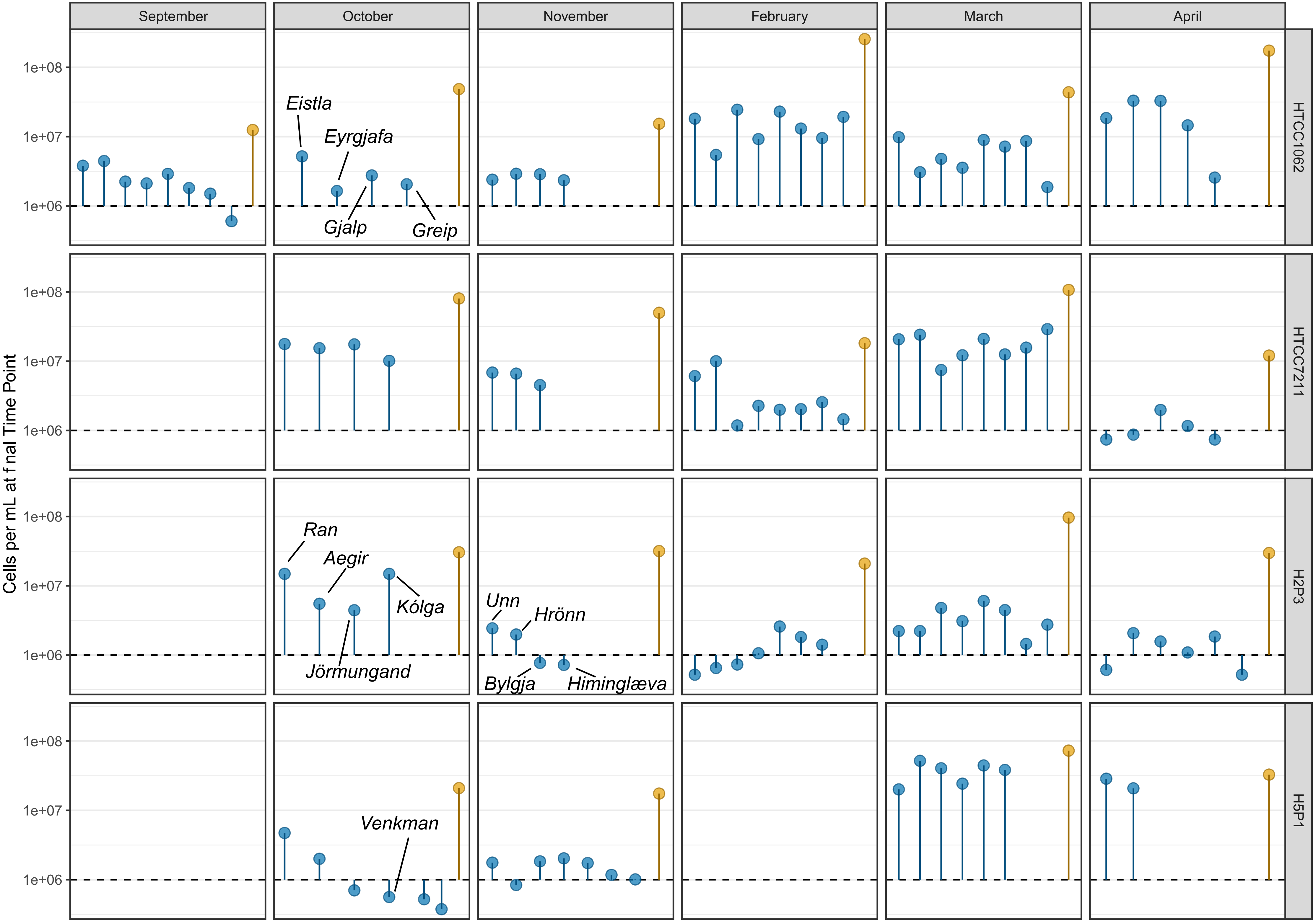

September
October
November
February
March
April
HTCC1062
HTCC7211
H2P3
H5P1
Cells per mL at final Time Point
1e+08
1e+07
1e+06
Eistla
Eyrgjafa
Gjalp
Greip
Ran
Aegir
Kólga
Jörmungand
Unn
Hrönn
Bylgja
Himinglæva
Venkman

51. Quast C, Pruesse E, Yilmaz P, Gerken J, Schweer T, Yarza P, et al. The SILVA ribosomal RNA gene database project: improved data processing and web-based tools. *Nucleic Acids Res* 2013; **41**: D590–6.

52. Vergin KL, Beszteri B, Monier A, Thrash JC, Temperton B, Treusch AH, et al. High-resolution SAR11 ecotype dynamics at the Bermuda Atlantic Time-series Study site by phylogenetic placement of pyrosequences. *ISME J* 2013; **7**: 1322–1332.

53. Eren AM, Borisy GG, Huse SM, Mark Welch JL. Oligotyping analysis of the human oral microbiome. *Proc Natl Acad Sci U S A* 2014; **111**: E2875–84.

54. Wickham H. ggplot2: Elegant Graphics for Data Analysis. 2016. Springer-Verlag, New York.

### Supplementary Materials List

Supplementary Fig. 1 Maximum likelihood trees (100,000 generations, sampled every 10 generations, first 5000 trees burn-in) of 16S rRNA genes of **(A)** Known SAR11 strains and new isolate H2P3α; **(B)** New OM43 isolates and other related members of Methylophilaceae. Trees in A and B were rooted using subclade III and the *Methylophilus*, respectively.

Supplementary Fig. 2 Cytograms of infected vs. uninfected SAR11 warm-water ecotype HTCC7211 cultures to isolate phages. Events within the acquisition range ("populations") of no-virus controls range from 72.7% - 84.3% of total events, whereas infected cultures range from 27.7% - 78.6%. Note the increase in low fluorescence (FITC-A) events for phage cultures, which were used to identify infections in cultures.

Supplementary Fig. 3 Cytograms of infected vs. uninfected SAR11 cold-water ecotype HTCC1062 cultures to isolate phages. Events within the acquisition range ("populations") of no-virus controls range from 72.7% - 80.5% of total events, whereas infected cultures range from 40.7% - 81.8%. Note the increase in low fluorescence (FITC-A) events for phage cultures, which were used to identify infections in cultures.

Supplementary Fig. 4 Cytograms of infected vs. uninfected new SAR11 isolate H2P3α cultures to isolate phages. Events within the acquisition range ("populations") of no-virus controls range from 72.8% - 76.1% of total events, whereas infected cultures range from 16.5% - 72.0%. Note the increase in low fluorescence (FITC-A) events for phage cultures, which were used to identify infections in cultures.

Supplementary Fig. 5 Cytograms of infected vs. uninfected new OM43 isolate H5P1 cultures to isolate phages. Events within the acquisition range ("populations") of no-virus controls range from 46.0% - 66.5% of total events, whereas infected cultures range from 15.7% - 56.8%. Note the increase in low fluorescence (FITC-A) events for phage cultures, which were used to identify infections in cultures. There was no clear discrimination between noise and the population peaks of the cytograms for infected cultures in March and April, therefore calculated percentages are likely influenced by application of gating parameters defined in other samples.

Supplementary Fig. 6 Distribution of pairwise average nucleotide identity (ANI) of all phages included in this study. The secondary peak at ANI >95% supports current use of this value as a cut-off for defining viral populations.

Supplementary Fig. 7 Unrooted maximum-likelihood tree of TerL genes found in *Pelagibacter* phages, using *Bordetella* phage LK3 as outgroup. Distinct branches were highlighted and labelled according to their respective hypergeometric cluster and viral family. Sampling environment of metagenomes used to identify pelagiphage-like contigs are marked: Epipelagic (TT-EPI mesopelagic (TT-MES), bathypelagic (BATHY), Southern Ocean/ Antarctica (ANT), Arctic Ocean (ARC). Branch support values are available within the newick file at https://github.com/ViralPirates/viral-dte.

Supplementary Fig. 8 Shared protein cluster network content of known phage isolates [33, 34] and our viral isolates against the NCBI Bacterial and Archaeal Viral RefSeq v85 with ICTV + NCBI taxonomy database (VConTACT2, default settings); different colours represent the ICTV-recognised genera assignments.

Supplementary Fig. 9 Relative abundance of SAR11 clade in Western English Channel at the time when samples were collected for viral isolation. (A) Total relative contribution. (B) Clade composition of the SAR11 fraction. The stacked barplots depict the relative contribution of each of the four clades to the total SAR11 fraction. NA: Not assigned to any of the four clades.

Supplementary Fig. 10 Growth curves of *Pelagibacter sp.* H2P3α, based on eight replicates in 125 mL flask of ASM1 medium, grown at different temperatures. Error bars represent ± std. error.

Supplementary Fig. 11 **(A)** Fluorescence and **(B)** Temperature at Western English Channel sampling station L4 over the course of the study. Grey dashed lines represent samples for viral isolation. Sampling missed phytoplankton blooms, potentially explaining the lower success of isolating viruses for OM43 strains.

Supplementary Fig. 12 Gene map of pelagiphages in Clusters A and B arranged inner to outer: *Eistla* (EXEVC025P), *Ran* (EXEVC014P), *Bylgja* (EXEVC010P) and *Eyrgjafa* (EXEVC018P). Solid lines indicate orthologous genes, dashed lines indicate genes of similar function.

Supplementary Fig. 13 Final cell densities of infected (blue) vs. uninfected (orange) hosts for 115 of the 117 viruses isolated in this study (two viruses isolated on OM43 D12P1 in July omitted). Dashed line at $10^6$ cells·mL$^{-1}$ represents host density at $T_0$. Points below this line indicate viruses reduced host density below the inoculum density. Named points refer to genomes sequenced in this study.

Supplementary Table 1 Pairwise average nucleotide identity of the 16S rRNA genes of OM43 isolates from this study.

Supplementary Table 2 Metadata associated with water samples taken for viral isolation.

Supplementary Table 3 Number of rounds of enrichment required for each water sample vs. target host combination before infection could be observed by flow cytometry.

Supplementary Table 4 Matrix of pairwise average nucleotide identity (ANI) for viruses from [33, 34] and this study.

Supplementary Table 5 General data and descriptions of sequenced viral isolates.

Supplementary Video 1 Time series of an infection of *Pelagibacter ubique* HTCC1062 showing how host cell density achieves a steady state in infected cultures that is lower than that of uninfected cultures. Associated cytograms are shown to demonstrate shifts in fluorescence associated with viral infection.
